# Comparative Analysis of Dimension Reduction Methods for Cytometry by Time-of-Flight Data

**DOI:** 10.1101/2022.04.26.489549

**Authors:** Kaiwen Wang, Yuqiu Yang, Fangjiang Wu, Bing Song, Xinlei Wang, Tao Wang

**Author notes:** These authors jointly supervised this work (Xinlei Wang, Tao Wang).

## Abstract

While experimental and informatic techniques around single cell sequencing (scRNA-seq) are advanced, research around mass cytometry (CyTOF) data analysis has severely lagged behind. CyTOF data are dramatically different from scRNA-seq data in many aspects. This calls for the evaluation and development of computational methods specific for CyTOF data. Dimension reduction (DR) is one of the critical steps of single cell data analysis. Here, we benchmark the performances of 21 DR methods on 110 real and 425 synthetic CyTOF samples. We find that less well-known methods like SAUCIE, SQuaD-MDS, and scvis are the overall best performers. In particular, SAUCIE and scvis are well balanced, SQuaD-MDS excels at structure preservation, whereas UMAP has great downstream analysis performance. We also find that t-SNE (along with SQuad-MDS/t-SNE Hybrid) possesses the best local structure preservation. Nevertheless, there is a high level of complementarity between these tools, so the choice of method should depend on the underlying data structure and the analytical needs.

## INTRODUCTION

Recently developed single-cell profiling technologies hold the promise to provide critical insights into the cellular heterogeneity in tissues of various biological conditions, developmental trajectories of single cells, and how cells communicate with each other. Researchers in the field of genomics and proteomics have separately developed single cell profiling technologies, mainly single cell RNA-sequencing (scRNA-seq) and flow cytometry, and their variants. Mass cytometry^1–4^, or CyTOF (Fluidigm), is a recent variation of flow cytometry, in which antibodies are labeled with heavy metal ion tags rather than fluorochromes. CyTOF captures much higher numbers of protein markers, compared with traditional flow cytometry, and has minimum spill- over effect, unlike regular flow cytometry^5^. Compared with scRNA-seq, CyTOF profiles the proteomics makeup of the single cells and is more relevant for understanding clinical phenotypes than scRNA-seq. CyTOF can additionally capture post-translational modifications^6^, which is beyond the reach of scRNA-seq. CyTOF is becoming increasingly popular, at an exponential rate similar to scRNA-seq (**Supplementary Fig. 1**).

There are key differences between CyTOF and scRNA-seq data, which makes it questionable to directly translate scRNA-seq analytical methods for use on CyTOF data. For example, scRNA-seq usually profiles several thousand genes’ RNA expression for several thousand cells, while CyTOF can profile up to 120 proteins’ expression for potentially 10 to 100 times more cells. The large number of cells enable CyTOF to capture the rare populations that may be missed by scRNA-seq data. CyTOF data uniquely suffers from time-dependent signal drift during the generation of data by the CyTOF machine. But CyTOF is mostly free from the drop-out issue that is commonly seen in scRNA-seq data^7–9^. Furthermore, while scRNA-seq data are integer count observations, CyTOF data are often regarded as continuous observations (despite that raw CyTOF data are still counts, but have a much larger range than those of scRNA-seq and will be pre-processed in various steps, which loses their discrete count nature). On the other hand, more and more studies are now trying to generate matched CyTOF and scRNA-seq data for the single cells from the same research subjects^10–12^. These two data types are of different nature, and thus are highly complementary with each other. Therefore, analysis processes should verify that findings and conclusions generated from combining CyTOF and scRNA-seq data are consistent/concordant. Unfortunately, these issues have not been addressed thoroughly by researchers, resulting in difficulties for valid interpretation of CyTOF data.

The first step of single cell data analysis is data exploration, which is usually achieved through Dimension Reduction (DR), and followed by visualization, clustering, cell type assignment, differential expression analyses, *etc*. Proper DR is fundamental to all downstream analyses. For example, if DR incorrectly places some cells in the wrong place in the reduced space, these cells could be mistakenly labeled as a “new” differentiation stage of another irrelevant cell type instead of their actual cell types. Traditional linear DR algorithms such as Principal Component Analyses have existed for decades. With the rise of scRNA-seq, numerous new DR algorithms, such as tSNE^13^, UMAP^14^, SAUCIE^15^, ZIFA^16^, PHATE^17^, scvis^18^, and Diffusion map^19^, have been developed or applied to capture the complicated non-linear relationships in the high-dimensional data. However, it’s unclear which of these DR methods is the best for CyTOF. Researchers have focused on benchmarking DR methods for scRNA-seq data in previous works^20, 21^. However, the best DR methods for scRNA-seq data may not necessarily extrapolate to CyTOF.

To fill in this void, we review 24 two-dimensional DR methods (**Supplementary Data 1**) and systematically compare the performances of 21 DR methods for CyTOF, based on 110 real and 425 simulated samples. To the best of our knowledge, this is the first study that has comprehensively reviewed and benchmarked DR methods for CyTOF data. We also include 10 Imaging CyTOF/mass cytometry (IMC) samples^22^, which is an expansion of mass cytometry that enables the capturing of the spatial information of the cells (**Supplementary Fig. 2**). Overall, our results rebut the common thinking in the field that tSNE and UMAP, the top performers for scRNA-seq data, are also optimal for CyTOF data. We find significant complementarity between DR tools, and we postulate that the choice of method should depend on the underlying data structure and the analytical needs. Our evaluation highlights some challenges for current methods, and our evaluation strategy can be useful to spearhead the development of new tools that effectively perform DR for CyTOF data.

## RESULTS

### Overall study design

We collected a total of 24 DR methods published by various researchers (**Supplementary Data 1**, and **Fig. 1a**), including those general purpose DR methods and those developed specifically for single cell-type of data. Among these methods, 21 were practically executable. We benchmarked them on a total of 110 real CyTOF samples from 11 studies (**Fig. 1b**, and **Supplementary Data 2**), including both peripheral blood and solid tissues, and also 425 simulated CyTOF samples of diverse characteristics, using our *Cytomulate* algorithm^23^ (**Fig. 1c**). Real data samples range from 5,024 to 604,081 cells and from 13 to 41 protein channels. For simulation datasets, we systematically varied the parameters, resulting in cell counts from 10,000 to 600,000 and protein channels from 30 to 60. Following the recommendations by its authors, scvis was benchmarked using subsampled CyTOF data for training the neural network. However, we also chose to test scvis (Full) by using full CyTOF samples.

**Fig. 1.**
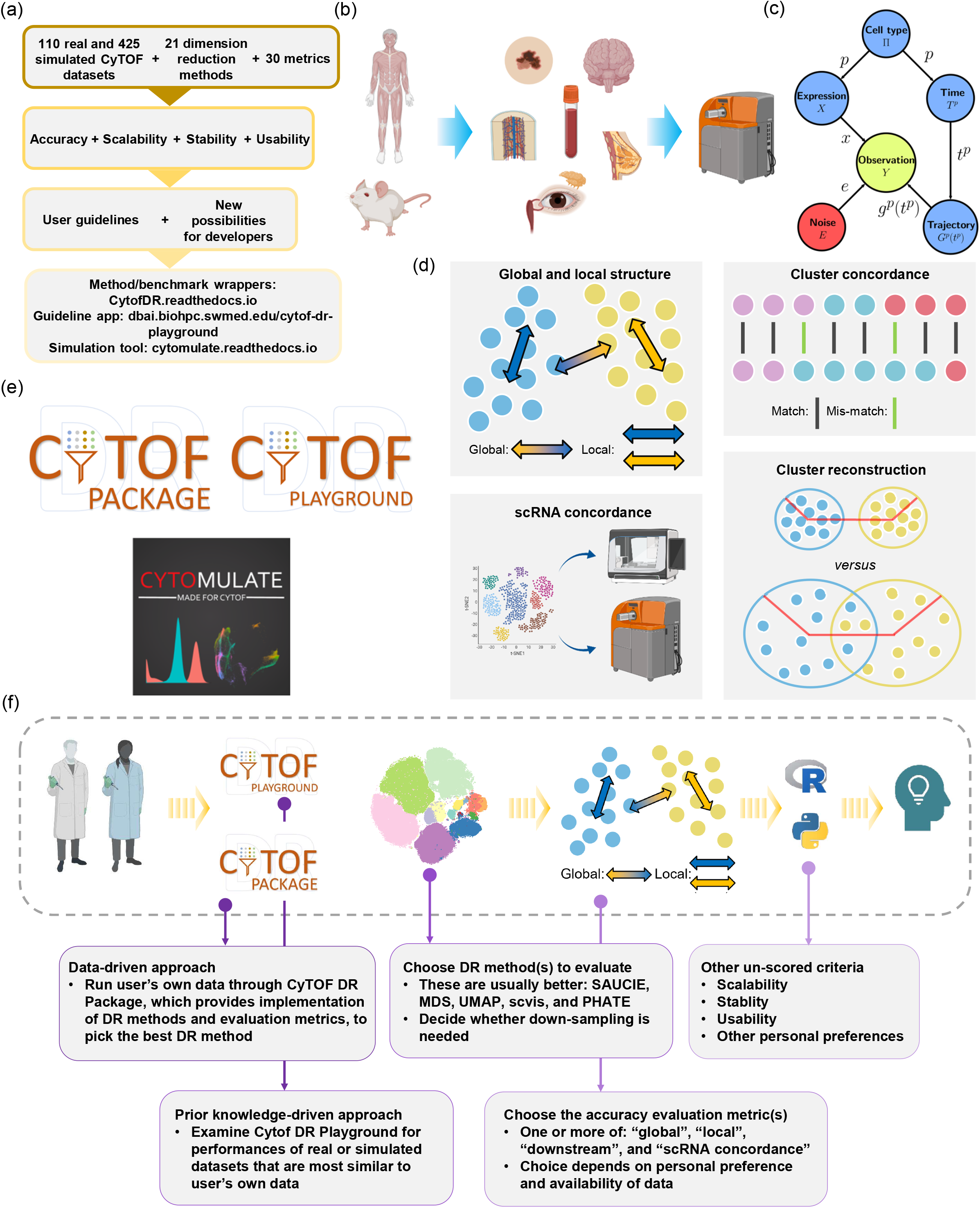
Overview of several key aspects of this benchmark study. (a) The datasets, DR methods, validation metrics, and deliverables of this study. (b) Synopsis of the real CyTOF datasets used for evaluation in this study. (c) The simulation algorithm for CyTOF data. (d) Diagrams explaining several of the most important validation metrics in the accuracy category. (e) Deliverables of this study, including, a complete set of guidelines for choosing DR methods based on data characteristics and user preferences (CyTOF DR Playground), our streamlined pipeline for implementation of the DR methods and evaluation metrics (CyTOF DR Package), and the *Cytomulate* tool. (f) A complete set of guidelines for users of DR methods to choose the best method for their CyTOF data.

For scoring the accuracy of the DR methods, we chose a total of 16 metrics in 4 main categories, characterizing different aspects of the performances of the DR methods. These 4 main categories are (1) global structure preservation, (2) local structure preservation, (3) downstream analysis performance, and (4) concordance of DR results with matched scRNA data. We provided a detailed description of these metrics in **Supplementary Data 3** and the **method section**. Some of the most important metrics were visually demonstrated in **Fig. 1d**. In our study, the DR methods were assessed and ranked mainly by these metrics focusing on the accuracy. But we also assessed and reported their scalability with respect to the number of cells and protein markers, stability of the DR after re-sampling the datasets and also parameter tuning; and the usability of the tools in terms of software and documentation (**Fig. 1a**, and **Supplementary Data 3**). More details about each assessment criterion will be given later. Employment of a similar set of metrics has also been adopted in other bioinformatics benchmark studies, such as Huang *et al*^21^ and Saelens *et al*^24^.

To operationalize our benchmarking framework, we gathered both publicly available and in-house real datasets (**Supplementary Data 2**), which provided a gold standard in terms of biological implications, and also generated synthetic datasets, which offer the most comprehensive and controllable coverage of different characteristics of CyTOF data. These real datasets come from a variety of organisms, dynamic processes, and types of trajectory topologies. In particular, while CyTOF has been applied for peripheral blood by immunologists mostly, we included CyTOF data from solid tissue experiments. In our real datasets, we also included 10 Imaging CyTOF datasets from one cohort created on breast cancer tissues (the BC cohort of **Supplementary Data 2**). For simulation of CyTOF data, we developed *Cytomulate*^23^ (**method section**), which to the best of our knowledge is the first formal simulation tool that is able to well mimic the behaviors of CyTOF data for general-purpose usage. Prior works, like LAMBDA^25^, have created specialized procedures to generate simulated CyTOF data in the context of model-based clustering. Such *ad hoc* simulation procedures cannot be used to test methods developed for other purposes (e.g. DR), whereas *Cytomulate* aims to support all facets of CyTOF research and methodological development with flexibility.

We also developed the CyTOF DR Playground webserver to help future users of DR methods visualize our benchmark results of these 535 CyTOF datasets, and choose the optimal DR methods for their own datasets (**Fig. 1e**). This app allows users to query the results of this evaluation study, including filtering of datasets and customized selection of evaluation metrics for final ranking. We also offer our unified implementation pipeline for homogenized input/output of the DR methods and the benchmark metrics (**Fig. 1e**, CyTOF DR Package), so that future users can easily execute DR and the benchmark metrics for their own data. Users can refer to CyTOF DR Package and CyTOF DR Playground to decide the best DR method for their datasets of interest (**Fig. 1f** and **Supplementary Note 1**). The user can either take a data-driven approach to run their dataset through all chosen DR methods based on their chosen evaluation metrics using CyTOF DR Package. Or the user can choose a prior-knowledge driven approach to query the best DR method from CyTOF DR Playground and make decisions based on similarity of the characteristics of their CyTOF dataset with those of the datasets we evaluated in this study. As another important deliverable of our work, we also packaged our CyTOF data simulation algorithm into *Cytomulate*, and shared it through cytomulate.readthedocs.io (**Fig. 1e**) for researchers to perform other CyTOF-related research.

### Accuracy of DR methods

We found that the top methods in terms of accuracy depend highly on the real CyTOF datasets and their characteristics, such as their tissue and disease types, suggesting the lack of a uniformly most accurate method which fits the need for all users with any CyTOF datasets (**Fig. 2** and **Fig. 3a**; the meanings of the accuracy metric abbreviations provided in **Supplementary Data 3**). The differences of the top DR methods in terms of ranking is particularly obvious across different cohorts. UMAP is one of the best DR methods for scRNA-seq data, and UMAP also performed decently well on CyTOF data (**Fig. 3a**). But it is somewhat surprising that less well-known or less used methods, such as SAUCIE, SQuaD-MDS Hybrid, and scvis, out-performed UMAP on many datasets (Wilcoxon Signed Rank Tests on overall accuracy: SAUCIE-UMAP *FDR* < *2*.*07* × *10*^-9^, SQuaD-MDS Hybrid-UMAP *FDR* < *10*^-15^, and scvis-UMAP *FDR* ≈ *0*.*0084*). tSNE and DiffMap are also popular scRNA-seq DR methods. On CyTOF data, tSNE’s performance is good with particularly strong performance in the Breast cancer (BC) Imaging CyTOF cohort, but lags slightly behind top methods in others. DiffMap is one of the bottom-ranking methods for CyTOF data. On the Covid CyTOF datasets, PHATE is surprisingly one of the leading methods along with UMAP.

**Fig. 2.**
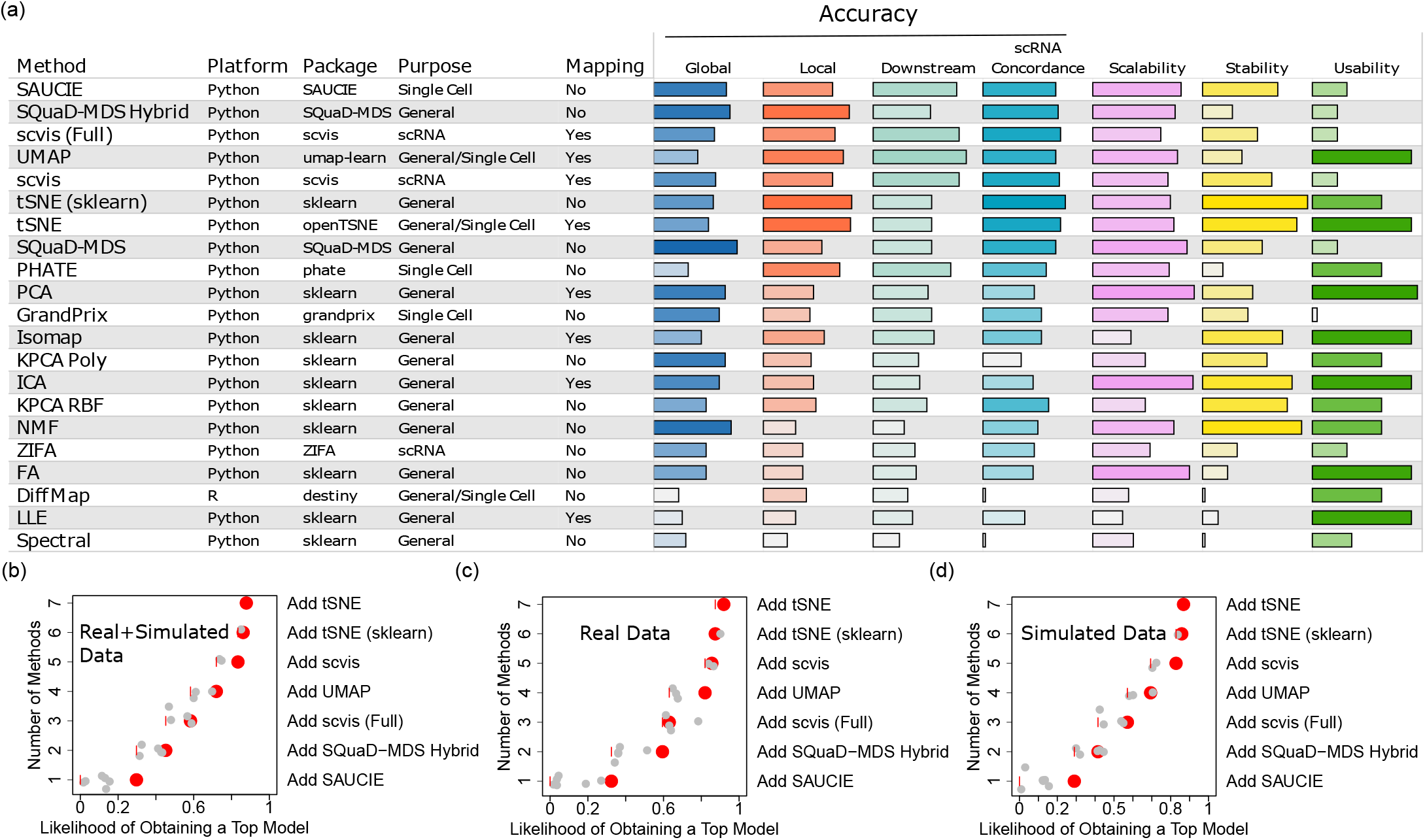
Overview of the benchmark study. (a) The DR methods reviewed in this study, their basic characteristics described in text, and their accuracy performances in terms of accuracy, scalability, stability, and utility. Results for the real and the simulation data were averaged for each of the four sub-categories. The DR methods were ranked based on the overall accuracy, averaging across all four sub-categories. All bars shown are calculated using ranks. Darker shaded bars within each column correspond to the better performance and therefore longer bars. Different colors are used to distinguish between different categories. (b-d) Complementarity of the DR methods, evaluated on the real+simulated CyTOF datasets (b), the real CyTOF datasets (c), and the simulated datasets (d). We define complementarity as the likelihood of obtaining a top-performing method for a given dataset by choosing a specific DR method. The short vertical red lines represent the baseline. The red points represent the resulting likelihood of obtaining a top model by adding the best method that has not been previously added. The gray points show the resulting likelihoods if other methods are chosen instead of the remaining best one. Small random noises are added to the y-axis to differentiate the gray points. Source data are provided as a Source Data file.

**Fig. 3.**
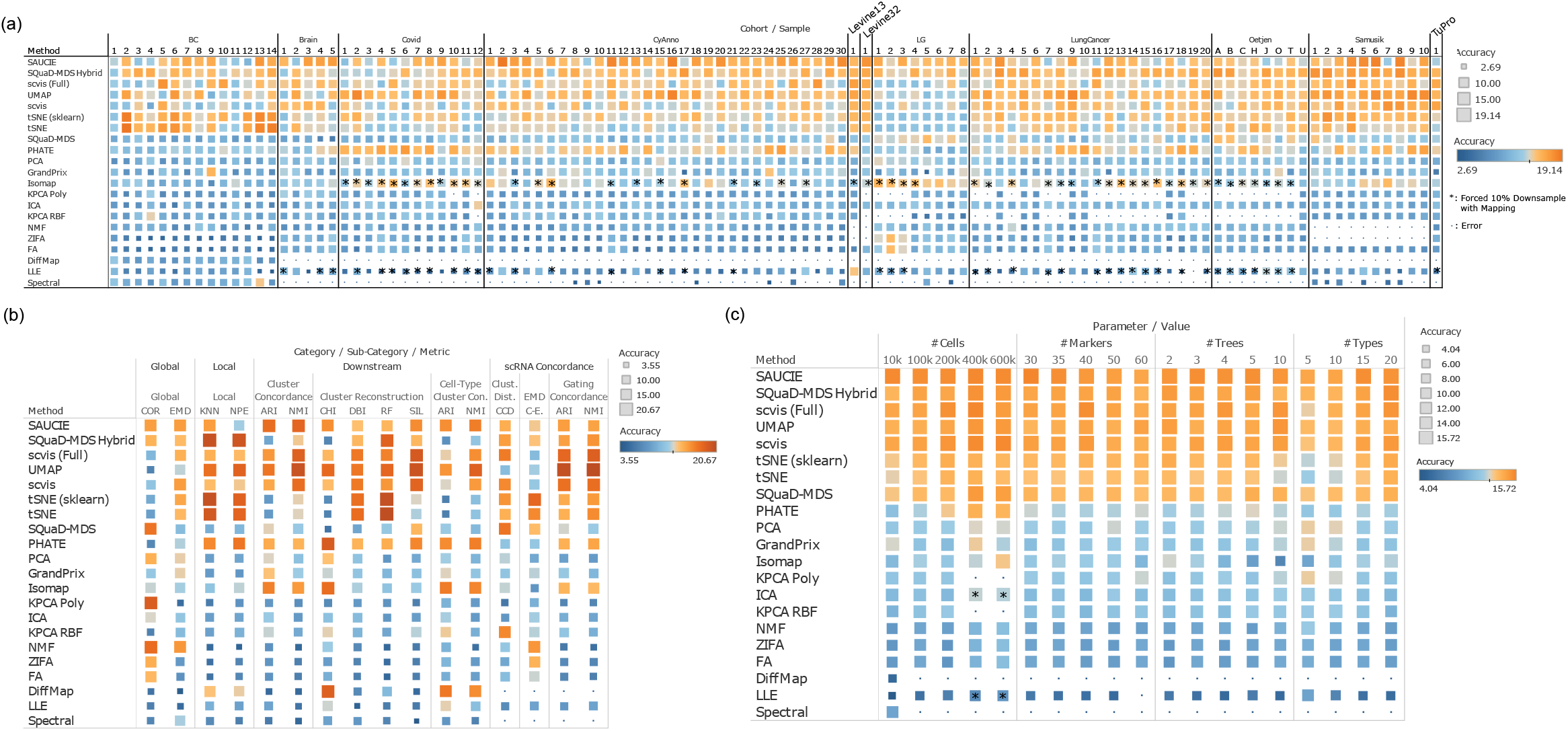
Detailed results on the accuracy performances of the DR methods. (a) The accuracy performances of the DR methods on all real CyTOF datasets, grouped by the 11 cohorts. Orange color indicates better performance, while the blue color indicates worse performance. All sub-metrics of the accuracy criterion were averaged. “*” indicates that forced 10% down-sampling was performed for training in the case that the methods aborted because of sample size. “scvis” is always performed on 10% down-samples per authors’ recommendation. Small blue dot indicates error. (b) The accuracy performances of the DR methods for each accuracy category. Results for all real CyTOF datasets were averaged, and the results for each detailed sub-metric of accuracy was shown. Orange color indicates better performance, while the blue color indicates worse performance. (c) The accuracy performances on the DR methods on the simulated CyTOF data. The performances were visualized as a function of the characteristics of the CyTOF datasets. Orange color indicates better performance, while the blue color indicates worse performance. All sub-metrics of the accuracy criterion were averaged. “*” Indicates that 10% down-sampling was performed. Source data are provided as a Source Data file.

We also examined the different categories of the accuracy metrics, and identified even stronger complementarity between the DR methods (**Fig. 3b**). For the purpose of this analysis, the performances of the metrics on all real CyTOF samples were averaged. We found that SAUCIE tends to perform generally well on most categories. SQuaD-MDS is also good overall except for a few submetrics in the downstream analysis performance. Scvis and UMAP are better at downstream analysis needs involving clustering and cell type assignment, and achieving concordance with the matched scRNA-seq data *via* gating concordance. tSNE and SQuad-MDS, the latter of which is based on both MDS and tSNE, are unequivocally the strongest in local performance and overall concordance with matched scRNA-seq datasets. PHATE also has decent performances on many metrics, but not including global structure preservation, which is rather poor. Further, there have been some hot debates regarding the global and local structure preserving capabilities of tSNE and UMAP^26, 27^. We found, in the context of CyTOF data, tSNE is better at both global and local structure preservation than UMAP (Wilcoxon Signed Rank Test *p* ≈ *1*.*23* × *10*^-6^ for global and *p* ≈ *2*.*43* × *10*^-5^ for local), but they are both inferior to SAUCIE in global structure preservation (Wilcoxon Signed Rank Test *p* < *10*^-10^ for both local and global comparisons between SAUCIE and UMAP and tSNE). This seems to conform to the results of Huang *et al*^21^, who benchmarked several DR methods in scRNA-seq data, and also found that the global preservation performance of UMAP and tSNE is suboptimal. Finally, tSNE’s downstream performance is mixed: while its RF and DBI metrics are favorable as compared to other methods, it lags behind in terms of other downstream metrics. This mixed result suggests that while local structure as a strength for tSNE is important for some downstream tasks such as classification, other characteristics of the embeddings play a role as well.

To more accurately evaluate how the performances of the DR methods vary as a function of the characteristics of the CyTOF datasets, we generated 425 simulated CyTOF datasets with the *Cytomulate* tool. We varied the numbers of cells, protein markers, independent cellular trajectories (#trees), and cell types in the simulated CyTOF datasets. We benchmarked the accuracy of the DR methods on these simulated data with the same criteria (**Fig. 3c**). The overall ranking of the DR methods in the simulation data is mostly consistent with their ranking in the real data (**Supplementary Fig. 3**). Most methods’ performances increase as a function of the number of cells, except for SAUCIE, which is mostly stable. Both tSNE variants and both scvis versions are sensitive to cell types as their performances are better with more cell types. Most methods’ performances are relatively stable with respect to the number of protein markers in the CyTOF data and the number of independent cellular differentiation trajectories. One exception is that both tSNE variants have decreased performances when the number of trajectories is large.

The variability of the top methods from different samples suggest complementarity between the DR methods (**Fig. 2b-d**). To quantify complementarity, we calculated the likelihood of obtaining a top DR result by using an increasing set of top DR methods. In real and simulated CyTOF data combined (**Fig. 2b**), we observed that using a single method (SAUCIE) for DR of CyTOF data can only generate “top” results ∼30% of the time. Adding a second method can guarantee a top result ∼45% of times, and top 4 methods yield 72%. Considering all top 7 methods together can lead to a close to 88% chance of obtaining a top result. Analysis on the real CyTOF data (**Fig. 2c**) and the simulated CyTOF data alone (**Fig. 2d**) also revealed strong complementarity between these different DR methods. Consequently, our results provide good evidence that users should not adopt a one-and-done approach. Instead, it is advisable to fit a number of top methods to not only maximize their likelihood of reproducing the original data more accurately, but also make decisions based on the advantages of each method tested (further summarized in the **Discussion** section) as an ensemble approach.

### Visual assessment of DR accuracy

Data visualization is an important downstream task contingent on DR. We embedded the Levine32 cohort with five top and popular DR methods in **Fig 4a**. The scatter plot of cells in the DR space are shown along with cell types and select metrics in terms of ranks. In **Fig 4a**, we chose to display three accuracy metrics for comparison purposes: COR, KNN, and ARI, to represent the Global, Local, and Downstream categories (more details in the **Materials and Methods** section and **Supplementary Data 3**). The top methods, in general, place the same or related types of cells in close proximity (e.g. CD8^+^ T Cells and CD4^+^ T Cells), which is a desirable characteristic overall. However, when examined in greater details, SQuaD-MDS Hybrid and tSNE render some of the cell types mixed and hard to distinguish from one another. These two methods have superb local structure preservation, which may help explain their placement of small clusters and the tight rendering of clusters, but their downstream performance, which relies on distinguishing between clusters and cell types, is not as impressive.

**Fig. 4.**
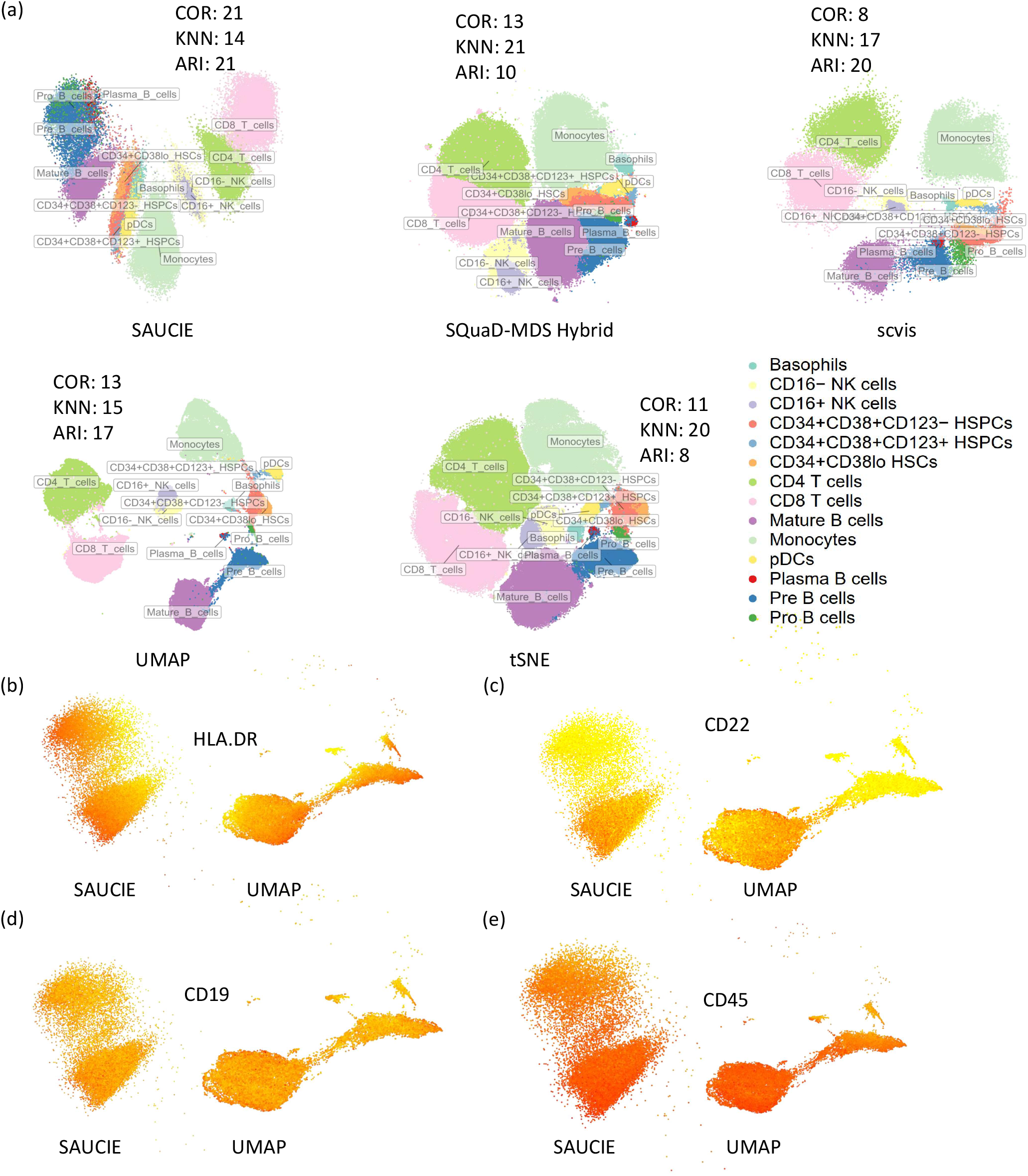
Visualizing DR results for the Levine32 dataset. (a) Visualization of the DR results of several top and popular methods on the Levine32 dataset, with cells colored by their cell types. The COR, KNN and ARI accuracy metrics, for each method, were also labeled beside each plot. (b-e) Visualization of the DR results from SAUCIE and UMAP, for the B cells only. In each panel, the cells were colored by the expression of HLA.DR (b), CD22 (c), CD19 (d), and CD45 (e). Red refers to higher expression and yellow refers to lower expression. Source data are provided as a Source Data file.

In light of its popularity and good downstream performance across many datasets, UMAP was further compared with SAUCIE by examining the different stages of B cells and several key marker genes in this dataset. In **Fig, 4b**, the expression level of HLA.DR is highlighted in both embeddings for the B cells. Previous research has shown that B cells develop in many stages and have more of a continuous gradient in cellular developmental progress, rather than discrete stages. HLA.DR (as are other MHC class II genes) is an important marker of the progression of B cells through these different stages^28, 29^. In the SAUCIE embedding, HLA.DR-high B cells are clustered together towards the left side, whereas UMAP placed these cells at three distinct places pointing to different directions. Next, we examined CD22, which is a surface molecule expressed early during the ontogeny of B cells^30^. We found that B cells with high CD22 expressions are clustered towards the bottom cluster in the SAUCIE embedding. However, these cells are clustered at the outer rim of one of the clusters of cells in the UMAP embedding, which is counterintuitive (**Fig. 4c**). A similar pattern could be observed for CD19 (**Fig. 4d**). Finally, we examined CD45, which is also an important marker of B cell development^31^. In the left panel of **Fig. 4e** with SAUCIE embedding, the expression forms a continuous gradient from the bottom (high CD45) to the top (low CD45). However, in the UMAP embedding, the cluster that represents plasma B cells (there are two small clusters above the big cluster on the right, the first small cluster on the left) is separated from the rest of the B cells far away, but has cells of both very high and very low expression of CD45, which is not optimal. A more sensible embedding would cluster cells with high CD45 from this small cluster together with other cells with high CD45 in the big clusters. Alternatively, the small cluster as a whole should be connected with other cells, rather than forming a separate cluster.

B cell development forms a complicated continuous gradient, supported by the gradient changes of the marker genes we showed in **Fig 4b-e**. SAUCIE’s embedding is more concordant with the current literature showing a smooth continuum in the DR space. While UMAP’s embedding with various clusters and elongated shapes may be more visually attractive, SAUCIE seems to better capture the true underlying biological processes. In fact, this case study suggests that DR for CyTOF data is more nuanced than simple visual inspections. Rather, empirical accuracy metrics combined with biological insights and side-by-side comparisons as a holistic approach yield the most productive use of DR results.

We conducted a second case study on the BC cohort, which consists of imaging CyTOF samples. The various types of cells in the tumor microenvironment should form a spatial gradient^32^. For example, the tumor cells in one region could be more differentiated, and it could become more and more de-differentiated when the tumor cells grow towards another region. We calculated pairwise spatial distances between cells from the same clusters. Then, we examined the correlation between spatial distances between cells and the distances between these same cells in terms of protein expression. Our rationale is that if the DR space is more accurate, the clustering in the DR space would be more likely to yield clusters that are indeed more homogeneous in terms of cell types. Then the correlation (spatial *vs.* expression) would likely be higher, influenced more by the spatial gradient in protein expression. But if the DR space is not accurate, making the clustering also inaccurate, this correlation will be confused by the abrupt changes in protein expression due to the mixture of cell types, and the correlation will be likely lower. Indeed, we found that SAUCIE is overall the best, in terms of achieving the highest positive correlation overall (**Supplementary Note 1 Fig. 5**).

**Fig. 5.**
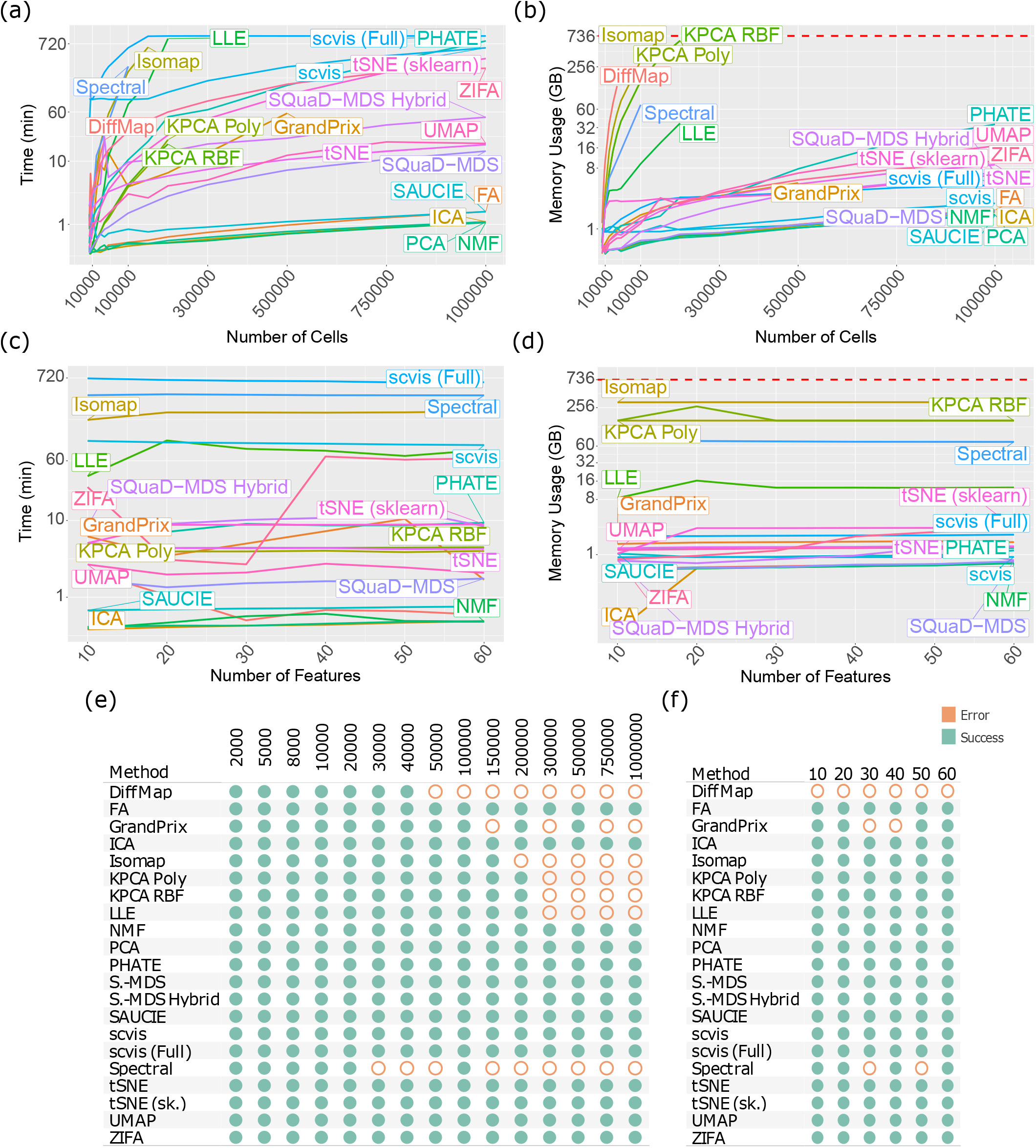
Detailed results on the scalability performances of the DR methods. The runtime (a,c) and memory usage consumption (b,d) of the DR methods. In (a) and (b), the DR methods were evaluated based on increasing numbers of cells. In (c) and (d), the DR methods were evaluated based on increasing numbers of protein features. (e,f) Whether the DR methods abort with error during the runtime/memory benchmarking analyses in each cell number group (e) and protein feature number group (f). Source data are provided as a Source Data file.

### Scalability of DR methods

As one of CyTOF’s unique advantages over other sequencing-based technologies, samples and cohorts often contain expressions of at least half a million cells, and the throughput of data is likely to further increase with development of the protocol and its rising popularity among researchers. Depending on the limitations of algorithms and their original purposes, not all methods are designed to handle such large datasets at this scale efficiently or at all. We provide a discussion on the theoretical efficiency and scalability of the top DR methods, with respect to the sample size dimensionality of CyTOF datasets, in **Supplementary Discussion 1**. To assess their scalability empirically, we also benchmarked each method with respect to their runtime and memory consumption using datasets up- and down-sampled from the Oetjen cohort^33^ (**Fig. 5ab**). Overall, runtime (**Fig. 5a**) and memory usage (**Fig. 5b**) seem to be highly correlated overall (Spearman’s *ρ* = *0*.*63*).

In terms of both runtime and memory usage, we found that the scalability of most methods was overall poor. Comparatively, SAUCIE, FA, ICA, PCA, NMF are most favorable in terms of runtime and memory usage in general. The most surprising result is SAUCIE’s efficiency, as neural networks typically are computationally intensive. Inspecting its architecture, we found that SAUCIE used a feedforward neural network with 7 layers: a 3-layer encoder, an embedding layer, and a 3-layer decode. This straightforward formulation ensures the efficiency of SAUCIE. UMAP, tSNE, SQuaD-MDS, and SQuaD-MDS Hybrid are in the next group of methods with slightly less efficient– but still reasonable – runtime and memory usage characteristics. The other methods are in the third tier with up to ∼100 times more consumption of runtime and memory.

The runtimes of these methods in our various experiments can be as much as 12 hours like scvis and the memory usage can be as large as 700GB. Scvis scales poorly with numbers of cells in terms of runtime. But its runtime plateaus with large numbers of cells at around 12 hours. Curiouclow, and very comparable with the first tier of methods (SAUCIE, *etc*).

Due to the scalability issue of many of these methods, subsampling the whole CyTOF datasets is sometimes necessary for some of these DR methods. For example, a number of methods, including Diffmap, Isomap, KPCA Poly/RBF, LLE, and Spectral, all tend to abort with increasing sample sizes. On the other hand, the original publications of some DR methods (e.g. scvis, UMAP, and PHATE) have demonstrated the robustness of their results with regard to subsampling. For the purpose of observing the overall cell state space of the cells (such as major cell populations and their relationships), subsampling, up to a certain extent, will probably not negatively impact the conclusions to be drawn. But all downstream analyses will have to be limited to the subsampled cells and rare cell populations may be missed in the down-sampled subset, which is the detrimental effect of subsampling. So whether to adopt subsampling or not also depends on the purpose of the analyses.

To address the scalability issue encountered while benchmarking the accuracy of DR methods, we took a two-pronged approach in our benchmark studies. For all accuracy benchmarks (e.g. results in **Fig 2** and **Fig. 3**), we did not allow subsampling unless a given method has a mapping function to produce an embedding with all cells. Under this setting, LLE and Isomap were the only methods that needed to downsample and had the capability of mapping new data: we used 10% sub-sampled data in select samples if they aborted on original samples, and the rest of the data were mapped onto the embedding for evaluation. For other inefficient methods without a mapping function, we scored these methods as NAs when they aborted, so that there would be no bias in our evaluation due to sample size differences. In addition, we conducted a separate benchmark to evaluate the performance of DR methods on all subsampled data. We found that the results are nearly identical to our main findings (**Supplementary Note 1 Fig. 7-8**). The exact sampling mechanism and evaluation procedure with subsampling are detailed in the **method section**. One exception to the aforementioned downsampling scheme is the inclusion of both scvis and scvis (Full) in our benchmarks. As recommended by its authors, we evaluated scvis on both the full CyTOF datasets and also down-sampled datasets with the mapping function.

Given the multitudes of implementations and optimizations of tSNE^13, 34–36^, we tested and observed that different tSNE implementations can have as much as a 10-fold difference for large samples (**Fig. 5a**). Specifically, our reference implementation with fast fourier transform (FFT) as implemented by openTSNE (named tSNE in this work unless otherwise noted), which is a relatively recent development, vastly outperforms the standard Sklearn version using the Barnes-Hut (BH) algorithm as sample sizes grow. Investigating further with more popular tSNE implementations, we again observed the overall runtime advantage of the FFT variants from “tSNE” and “tSNE (FIt-SNE Original)” (**Supplementary Fig. 4**). Surprisingly, the BH implementation from openTSNE outperforms all other implementations when sample sizes are small, while falling not far behind the FFT tier for large samples. With the large sample sizes of CyTOF data, it makes sense to always use FFT-based tSNE, but BH still offers some value for small samples and when the FFT variants are not easily accessible. The memory usage is more similar among these tSNE implementations, but still with several folds of differences.

On the other hand, we also benchmarked the scalability of the DR methods with respect to the dimensionality of the numbers of protein features. In the above experiments, we have a total of 34 protein markers in the CyTOF data (from the original Oetjen dataset), which is typical for most current CyTOF datasets. In consideration of the possibility that future iterations of the CyTOF technology could incorporate more protein markers, we up-sampled the Oetjen dataset to create up to 60 protein markers while fixing the cell number at 100,000. Then we performed the same benchmark study in terms of runtime (**Fig. 5c**) and memory usage (**Fig. 5d**). Our benchmark analyses show that most methods do not vary too much with respect to the number of protein markers in the CyTOF datasets, alleviating concerns for increased runtime/memory usage for future CyTOF data with more protein markers.

Finally, we also noticed, during our DR benchmark practices above, that some of these DR methods will abort with errors due to increasing dimensionality. To explicitly quantify the dimensionality threshold, we recorded whether the DR methods would abort while performing the scalability benchmarks of various dimensionalities. Specifically, the methods that produced errors in **Fig. 5a-b** and **Fig. 5c-d** were recorded in **Fig. 5e** and **Fig. 5f**, respectively. Since sample dimension and the scalability are of interest, we did not allow subsampling for any method here. Our analyses show that Diffmap, Isomap, KPCA Poly/RBF, LLE, and Spectral start to abort with errors with an increasing number of cells, whereas GrandPrix often fails to converge (**Fig. 5e**).

These behaviors are consistent with our observations in other benchmarks. With increasing numbers of protein markers, our analysis for error occurrences shows that Spectral and GrandPrix sporadically abort (**Fig. 5f**), and that Diffmap were unable to complete any of the runs (of this particular scalability benchmark dataset) due to their inherent sample size limitations.

Synthesizing all our scalability analysis results, it is obvious that the number of cells, but not number of protein markers, in the CyTOF data is the driving-factor in determining each method’s scalability and thus becomes an important consideration for end users. But unfortunately, CyTOF data inherently profiles at least an order of magnitude more cells than scRNA-seq data. Therefore, optimizing methods for data throughput should be one of the top priorities for researchers.

### Stability of DR methods

While the accuracy of DR methods reflects their ability to reproduce each sample faithfully in the embedding space, a crucial part of assessing their performance is also to ask whether each method can produce good results consistently. To test the stability of each method, we executed each method on 100 bootstrap samples of the Oetjen cohort’s Sample A (with the same tuning parameters), and calculated the stability of the DR methods, across different bootstrap datasets. Stability is defined as the average Earth Mover’s Distance (EMD) between pairwise distance measures from the embedding from the original sample and the bootstrap samples’ embeddings. This is a measure of overall structural difference globally using the difference in distribution, and the details of implementation are included in the **Methods** section. Smaller EMD values suggest that the DR method produces similar embedding under perturbations of the input. We found that most methods, except for PHATE and LLE, are relatively stable (**Fig. 6a**). The top methods that have the best stability performances are tSNE (sklearn) and NMF (average EMD shown in **Fig. 6a;** Wilcoxon Rank Sum test *FDR* ≈ *0*.*65* between tSNE and NMF, and *FDR* < *10*^-5^ for all other pairwise tests between tSNE and less stable methods; all pairwise tests *FDR* < *10*^-5^ between NMF and less stable methods). The second tier of methods includes tSNE, ICA, KPCA (RBF), and Isomap. The DR methods that have the best accuracy (**Fig. 2**) unfortunately do not possess the best stability performances. Among methods with decent accuracy, both tSNE variants are the most stable, which is not surprising because they both use PCA as initialization. Interestingly, SQuaD-MDS Hybrid (a tSNE-based method) is not as stable as other vanilla tSNE despite its accuracy advantage. SAUCIE, UMAP, and scvis’s stability performances are in the middle, suggesting future improvements to these methodologies are needed from method developers in this regard.

**Fig. 6.**
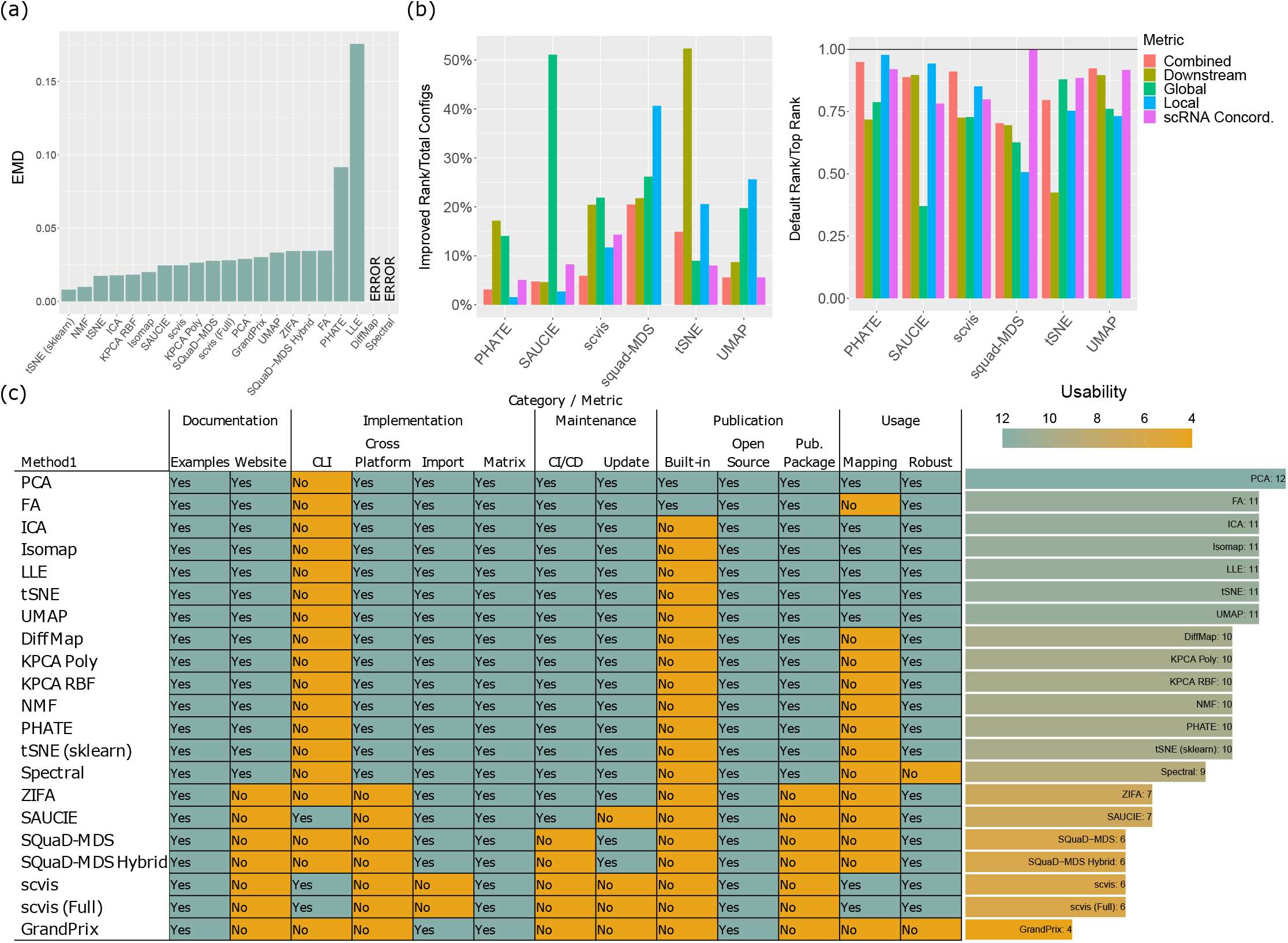
Detailed results on the stability and usability performances of the DR methods. (a) The stability of the DR methods. We calculated the stability of the DR methods, across different bootstrap datasets generated from the Oetjen cohort’s Sample A. (b) The impact of parameter tuning on the accuracy of the DR methods, in comparison with the default settings. In the left panel, we showed the optimal settings’ (among all tested settings, **Sup Table 4**) increases in ranks (over default settings’ ranks) divided by the total number of configurations for each method and each category. The right panel shows the relative performance of the default settings (with respect to the optimal settings), quantified by the ranks of the default settings divided by the maximum ranks achieved by the optimal settings for each method. (c) Usability scoring for the DR methods. All detailed usability criteria were displayed for all DR methods, in the format of a heatmap. Teal shading represents the presence of a feature in the heatmap and better overall usability in the bar graph, whereas orange shading indicates the lack of a feature and poorer usability. Source data are provided as a Source Data file.

Another important aspect of the DR methods related to the stability of computation is how much the results will vary as a function of their tuning parameters, given the same input data. Sometimes, it may be preferable that the DR results do not vary too much even given different parameters. But in other times, the data analytics practitioners may prefer that parameter tuning will lead to more diverse results given different parameters, so that they can choose a parameter setting that yields the optimal DR results. While we leave the choice of preference to the practitioners, we performed a study of parameter tuning. For several top DR performers (SQuaD- MDS, PHATE, SAUCIE, scvis, tSNE, and UMAP), we tuned their parameters, as is shown in **Supplementary Data 4**. We found that (**Fig. 6b**) all tuned methods benefit from careful selection of tuning parameters. tSNE, SQuaD-MDS, and scvis overall as well as the Global score of SAUCIE noticeably benefit the most in terms of percent rank improvement in accuracy, which is defined as the increase of ranks from the top configuration (as compared to the default) divided by the total number of configurations for each method. From the right panel of **Fig. 6b**, we observed that the default configuration oftentimes is a decent starting point, especially for UMAP and PHATE, and some accuracy categories of SAUCIE. While the default settings can often achieve above-average ranks in all settings tested, there is still room for improvement for every method. According to our analyses, we present the optimal configuration setting for each major category of accuracy in **Supplementary Data 4**, so that future users may choose to use these settings as the starting point for DR of their own CyTOF data. In general, we recommend using more training steps for SAUCIE while not changing any regularization coefficients. tSNE benefits from large perplexity, whereas using small minimum distance between points yields good results for UMAP. For SQuaD-MDS, using more iterations and a larger learning rate can offer improvements, especially Local performance. We do not recommend performing extensive parameter sweep for scvis, as the potential gain is small and also because scvis already performs well.

### Usability of DR methods

While not directly related to the accuracy of DR, it is also important to closely inspect each method’s implementation and evaluate methods based on both optimal user experience and quality of software. Based on our own experience working with each DR method, we employed a clearly-defined checklist to score each method’s usability, which includes documentation, automated testing and integration, regular update, cross platform compatibility, capability of accepting general matrix input, *etc* (**Supplementary Data 3** and **method section**). As shown in **Fig. 6c**, we found that popular methods and traditional methods, which were mostly implemented by well-known libraries such as sklearn, offer exceptional usability with not only class-leading documentation but also free of any quirks. However, working with some less well-known methods proved to be more difficult, especially when we were resolving installation issues, which stemmed from lack of software updates. Specifically, SAUCIE, scvis, and GrandPrix all depend on out-of-date tensorflow versions (1.x), which do not have support for recent versions of Python (3.8 and later). In this case, users have to resort to legacy software for these methods with questionable future support. Installation of ZIFA requires manual intervention but is relatively easy. Scvis is command line-only and thus cannot be flexibly invoked in R or Python, and the official implementation of DiffMap is in R only. Finally, we note that there are several methods that are capable of mapping new data onto existing DR embeddings (*e.g.* scvis, tSNE and UMAP), which could be very convenient in handling continuously generated data. Overall, some top methods like SAUCIE and scvis have very good performances for CyTOF data according to the accuracy criteria, but our work suggests that the field should consider improving their usability to maximize their benefit for CyTOF researchers.

## DISCUSSION

In this work, we present a comprehensive review and benchmark of popular and domain-specific DR methods across many different CyTOF datasets. The performances of the DR methods on the real and simulated CyTOF data are overall comparable, indicating the validity of our evaluation results as well as the simulation approach. Previous comparative works in the field of scRNA-seq have supported the notions that tSNE and UMAP are the top performers in general and even linear methods are well-suited for certain workflows^20, 37, 38^. Few attempts at tackling this issue for CyTOF data have been made and the field seems to think in general that the best methodologies for scRNA-seq data can be directly applied on CyTOF data. However, we found that for CyTOF data, SAUCIE, SQuaD-MDS Hybrid, and scvis (Full) are the overall top runners, proving the need for a CyTOF-specific benchmark study of DR methods. But at the same time, none of DR methods is perfect and there is a large degree of complementarity between them. On the other hand, our comparative results echo some of the previous conclusions regarding the performance of DR methods on scRNA-seq data. For example, previous studies have shown that tSNE performs better than UMAP in terms of local structure preservation on general and scRNA-seq data^21, 39, 40^. Our analyses on real and simulated CyTOF data demonstrate the same in general.

From the perspective of a practitioner working on CyTOF data, our results provide a set of good guidelines and conclusions. SAUCIE and scvis are all-rounders that perform admirably across all categories without major flaws, but SAUCIE is far more efficient than scvis. Thus, it is reasonable to use SAUCIE as part of the general workflow. The one caveat with SAUCIE is its questionable usability: while we would have easily recommended SAUCIE for rapid prototyping given its efficiency, the effort needed to set up SAUCIE renders it inferior to other efficient methods like PCA for this purpose. SQuaD-MDS Hybrid combines the local performance advantage of tSNE with the global performance of SQuaD–MDS, making it an overall excellent method for structural preservation. However, its downstream performance has some weaknesses, especially regarding cell-type cluster concordance. On the other hand, UMAP excels in downstream analyses with clustering workflows, but it sacrifices Global structure performance in return. We still recommend UMAP for tasks such as cell typing, clustering, and others that rely on tight and distinct clusters. Both tSNE variants have a unique set of strengths, but given the performance advantage of SQuaD-MDS Hybrid, we would recommend the latter instead for users who seek local accuracy. One key advantage of tSNE is its usability and cross-platform compatibility, which again highlights the inherent advantage of a more established method as compared to newer methods. PHATE is potentially good for some other downstream tasks that involve cell differentiation. On the occasions when Global structure is of concern, linear methods like NMF and PCA still make a strong case, but these methods are not as useful for downstream analyses. Overall, these results suggest that no one method dominates: users should be aware of each’s strengths and limitations. A holistic approach will be to start with a top and efficient method like SAUCIE or SQuaD-MDS Hybrid and then select some other methods for specific tasks and validation (*e.g.* UMAP for cell-typing).

Among these DR methods, we observed a polarized phenomenon that those popular DR methods usually have exceptional software and superior documentation, whereas some more accurate but less famous ones are severely lacking in this regard. The latter’s poor usability further aggravates the situation by discouraging practitioners from trying them out, which in turn demotivates the improvement and optimization of such methods. The current prevalent methods, such as UMAP and tSNE, turn out a result of a mix of outstanding performance, great usability, and good scalability. Their success, however, highlights the necessity to strike a balance between strong methodological research and user-oriented development. In light of these observations, we urge researchers to take end-user experiences and the increasing sample sizes of CyTOF into consideration and to develop competitive methods that shine on both theoretical performance and practical user experience. For example, SAUCIE is a neural network, which theoretically should have the capability of mapping new data points into an existing DR space. However, no such mapping functionality is provided in its software implementation yet.

Some researchers seem to analyze CyTOF in the same way as how flow cytometry data have been traditionally analyzed, using commercial software offering simplistic functionalities. They ignore the fact that CyTOF is a much more advanced technique that offers capturing of a lot more protein channels, but has minimum spill-over effects^5^. They also ignore all the sophisticated algorithms developed in recent years for cytometry data (flow cytometry, CyTOF, *etc*)^41^ and also for scRNA-seq data, which might be applied for analyzing CyTOF data. These new algorithms could reveal complicated biological insights that cannot be afforded by standardized commercial software. For example, it might be of interest to evaluate in the future whether the pseudotime inference algorithms that work well for scRNA-seq data are also suitable for pseudotime inference of CyTOF data^24^. On the other hand, we envision the DR benchmark results we obtained for CyTOF data in this study could be applicable for traditional flow cytometry data as well, including spectral flow cytometry^42^, which measures a number of protein markers that is on par with CyTOF, has higher cell throughput, but also still possesses higher spillover.

One curious observation we made is that, while SAUCIE and scvis are less well known than t-SNE and UMAP for DR of scRNA-seq data, these two methods turn out to be among the top tier of methods for DR of CyTOF data. This may in part be due to the fact that they are based on deep learning (DL) techniques, and their great performance for CyTOF data could be owing to the tremendous success of DL for handling big data (CyTOF captures much more number of cells than that for scRNA-seq data). Further, SAUCIE’s conventional autoencoder and scvis’s variational autoencoder both assume an underlying latent variable model with smaller dimension for features. CyTOF has much lower dimensionality, in terms of numbers of genes/proteins, which might have rendered this latent space inferred by SAUCIE and scvis more appropriate for capturing the true structure of CyTOF data. Beyond SAUCIE and scvis, many other methods (*e.g.* UMAP and tSNE) fall under the class of methods that are based on neighbors and optimizing distance measures, rather than DL approaches. Although the range of protein/gene features from CyTOF to scRNA-seq poses no computational concerns for these methods, the curse of dimensionality (*e.g.* the appropriateness of distance measures like Euclidean distance^43^) can still play a role in accuracy. Their overall lackluster Global performance compared to DL-based methods indicate the need for further theoretical considerations.

The fields of genomics and proteomics have been developing mostly independently. The ability to combine both approaches will be extremely powerful. There has been a growing number of studies generating single cell datasets with both scRNA-seq and CyTOF data^10–12, 44^. The creation of such matched datasets calls for the development of dedicated analytical methodologies to integrate these two types of data. First, one should ensure that the conclusions drawn from each of the proteomics and genomics modalities are overall consistent, as we reviewed in this work.

Second, robust and innovative statistical methods should be developed for the integrative analyses of CyTOF and scRNA-seq data, to maximize the potential of each technology. In this regard, CITE-seq^45^, which allows capture of the RNA transcriptomes and cell surface protein expression for the same cell, is unlikely to replace CyTOF (matched with regular scRNA-seq). CITE-seq can capture much less number of cells than CyTOF, and CITE-seq can only profile the expression of protein surface markers, while CyTOF is capable of capturing the expression of both intracellular and cell surface proteins^46^. CITE-seq also suffers from drop-outs in the protein expression measurements by the UMI counting approach^47, 48^, while CyTOF is relatively free from this caveat.

CyTOF possesses a much higher potential of being developed into a clinically applicable diagnostic/prognostic/predictive tool, compared with scRNA-seq, due to its cheaper per-cell cost, shorter workflow, and direct measurement of proteomics changes. Our study of the performances of DR methods on CyTOF data urges the field to rethink analytical strategies for CyTOF data and to develop state-of-the-art tools to address this key gap. This effort should ultimately lead to effective translational application of the CyTOF technology for personalized medicine.

## METHODS

### Benchmark environment

All compute- and memory-intensive benchmarking exercises were performed on server nodes with dual Intel Xeon E5-2695 v4 CPUs. Each node has 36 physical cores and was allocated 200 Gigabytes (GB) of RAM, except for all scalability benchmarks, where nodes with the same CPUs but 732 GB of RAM were assigned. All jobs were run in CentOS Linux and managed with Slurm with no GUI to ensure reproducibility. Each computation job was allowed to run for up to 7 days if necessary. Since no method exceeded this time limit, all errors and exceptions reported in the paper resulted from either out-of-memory errors or other unforeseen implementation reasons that we could not control. Statistical analyses, preprocessing, and other workflows for this paper were performed in R 4.0.4 with various consumer-grade hardware running Windows and Ubuntu.

### DR method implementation details

The benchmark pipeline was implemented mostly in Python (3.7 and 3.8) for its extensive support for DR methods and command-line operability. If available, Sklearn 0.24 implementations were used, as they are not only well maintained and but oftentimes also the default choice for Python users. For other DR methods, we utilized the implementations of the original authors if possible, with the exception of tSNE, which will be further discussed.

Additional Python DR packages used are: openTSNE 0.6, umap-learn 0.5, zifa 0.1, PHATE 1.0, scvis 0.1, SAUCIE (the version on GitHub https://github.com/KrishnaswamyLab/SAUCIE), GrandPrix 0.1, and SQuaD-MDS (the version on GitHub https://github.com/PierreLambert3/SQuaD-MDS-and-FItSNE-hybrid). One notable method to mention here is Diffmap since its reference implementation is available only in R (“destiny” 2.15.0 with R 3.6.3). As a convenient wrapper, our pipeline and our CyTOF DR Package offer interfaces for most methods. In our package and all analyses in this study, except for the parameter-tuning section, we used all default settings for tuning parameters to ensure fairness across all methods. But multicore optimizations (e.g. the “n_jobs” option in sklearn methods), if available, were enabled to utilize all cores on our server for efficiency.

“tSNE” in the main figures refers to the openTSNE implementation of FIt-SNE. Works such as Kobak *et al*^49^ have called for using FIt-SNE in the genomics field, because its computation efficiency is comparable with UMAP. So we choose this as our reference implementation. tSNE (FIt-SNE Original), which is the original authors’ implementation, technically offers faster speed and better memory efficiency, but it comes at a cost of usability with no discernable embedding improvement. Thus, openTSNE is a reasonable compromise that both improves upon efficiency as compared to BH and offers a user-friendly interface with good documentation. We also included “tSNE (sklearn)” in our main figures and benchmark study as it is one of the most popular choices with BH implementation. Other methods included in **Supplementary Fig. 4** serve as a scalability-only benchmark for users to understand the differences between the various choices.

For popular as well as top methods from our benchmark, we identified their tuning parameters and performed extensive parameter tuning. We list all the parameters in **Supplementary Data 4,** and for each method, we performed a grid search of all combinations. SAUCIE has the most parameters for both regularization (Lambda C and Lambda D) and optimization (Steps, Learning Rate, and Batch Size), yielding a staggering 1,125 total configurations. On the other hand, the authors of scvis have already tuned parameters and found that scvis is not sensitive to changes of settings. We thus validated their claims by tuning regularization coefficient, perplexity, and the number of neurons on layer 1 and 2.

### Clustering methods

For all CyTOF samples benchmarked in the study, we used FlowSOM^50^ for clustering in both the original space and the embedding space. FlowSOM has been the recommended algorithm for CyTOF because of its accuracy and efficiency^51, 52^. To ensure comparisons of DR methods are fair, we used the same original-space clustering labels for each sample during all benchmarks. FlowSOM requires an estimated number of clusters to be provided by the user. For cohorts with provided cell-typing information by their authors, we used the number of cell types as prior knowledge. For other cohorts, we determined the estimated number of clusters empirically using cluster variance and the elbow rule. Since FlowSOM also performs a meta-clustering step to consolidate clusters if necessary, we provide a larger number of clusters than the baseline obtained from the above methods (10-20 more clusters than the number of baseline clusters. The end results after consolidation are generally robust regardless of the choice of this number of clusters). Our approach aims to preserve the granularity of rare cell types while allowing the meta-clustering step to find the optimal clustering. All other settings were kept as default. Our study also includes a few matched scRNA-seq samples from the TuPro cohort and the Oetjen cohort. In these cases, we used the clustering algorithm provided by the Seurat^53–56^ pipeline as per the standard workflow for scRNA-seq data analyses. The number of clusters were determined automatically by the Seurat algorithm.

Admittedly, choosing different clustering algorithms and different tuning parameters could inevitably lead to a somewhat different benchmark result. This could happen no matter which algorithm and what set of tuning parameters we chose. Therefore, to ensure maximum fairness in good faith, we chose FlowSOM for CyTOF data as FlowSOM has been the recommended algorithm for CyTOF data analyses, and the clustering algorithm provided in Seurat, per the standard workflow for scRNA-seq data analyses. These two choices of clustering algorithms are likely the most common choices for CyTOF and scRNA-seq data users.

### Accuracy overview

Our main evaluation framework assesses the accuracy of each DR method’s embeddings. The framework consists of four major categories: global structure preservation (Global), local structure preservation (Local), downstream analysis performance (Downstream), and scRNA-seq concordance. Downstream and scRNA-seq Concordance have sub-categories to reflect different, equally-weighted aspects of the major categories. For each method, we utilized the original space expression data, the reduced space embeddings, clusterings before and after DR, assigned cell types, and, if available, the corresponding scRNA-seq data to execute each individual evaluation metric under major and sub-categories. The details and implementations of these methods are described later on in each of their own sections.

For each dataset, we first ranked all the methods based on each individual metric. Methods that aborted for whatever reason are ranked last for all metrics because they were unable to produce an embedding at all. In our weighting scheme for all metrics, we ensured that all major categories, all sub-categories, and all individual metrics within categories have the same weight (i.e. metrics at the same level are equally weighted). In practice, we first averaged the ranks of individual metrics under each sub-category if applicable. Then, the scores for each sub-category are averaged again as the scores for each major category. The Global and Local Category has no sub-category, and thus we directly averaged their respective metrics’ ranks. Finally, we averaged the scores of major categories to form a final accuracy score. The exact weight of each individual metric as part of its corresponding major category is detailed in the *Category Weight* column of **Supplementary Data 3**.

Given the rank-based nature of the accuracy scores, we compared and averaged scores across datasets from different cohorts, including both real and simulation data (**Fig. 2**). For accuracy-related figures (**Fig. 2**, **Fig. 3** and **Supplementary Fig. 3**), we ordered the methods according to the overall accuracy, combining all four sub-categories, in a descending order. Based on accuracy, we defined complementarity as the likelihood of obtaining a most accurate method if practitioners choose a subset of methods. In practice, we tallied the top-ranking methods for all real and simulation datasets and counted the number of times they were the top performer in terms of accuracy. Then, we calculated the observed proportion of each method being on top by dividing the number of times that each method ranked on top by the total number of datasets. A large likelihood means that the method is more often the best as compared to others, while 0 means that it was never the best in our benchmarks. We selected seven most accurate methods according to overall accuracy (**Fig. 2a**) and showed the increasing likelihood of obtaining a top method for a given dataset by adding each of the seven methods one by one in the order of overall accuracy (**Fig. 2b-d**).

### Accuracy - global structure preservation

This category aims to assess the degree to which each embedding preserves the structure of the original, high-dimensional data on a large scale. In practical terms, we aim to evaluate the relationship between each cell with not only its neighbors but also other cells that are vastly different from themselves. Metrics under this category are critical in helping practitioners decide whether the overall structure present in an embedding, such as the spatial relationship between a few clusters, is interpretable or significant.

Previous research^20^ has called for using pairwise distances in the original and dimension-reduced space to capture the relationship between all cells. The problem of implementing this for CyTOF datasets is that the memory complexity is *O*(*N*^2^), which is prohibitively expensive and practically impossible even using state-of-the-art servers. Thus, we implemented the Point-Cluster Distance (PCD) for all main benchmarks in this study. Instead of computing and storing the full pairwise distance vectors, we utilized the original space clustering of each sample to calculate the centroid of all clusters. Suppose there are *C* clusters with each cluster denoted as *c*. Then, we computed the Euclidean distance between each individual cell and each cluster centroid. This approach preserves both the granularity of each individual cell’s location and the overall structure of the data while also reducing the memory complexity to *O*(*N*) with an *N* × *C* matrix. Alternatively, another approach is to downsample the CyTOF datasets and to compute the accuracy scores within these subsets, which yields similar results as PCD. We included a further discussion on these two approaches in **Supplementary Note 1**.

Spearman’s Correlation: This metric measures the Spearman’s correlation of the original and dimension-reduced space using PCD. The PCD matrices from the original and DR space are flattened, and the correlation (*ρ*) coefficient is computed between the vectors. By utilizing ranked-based correlation, we no longer require the normality assumption on PCDs. The resulting *ρ* is an indication of how well the ranks are preserved, and a large, positive *ρ* is preferred, suggesting that cells that are relatively far apart in the original space remain relatively far apart and vice versa.

Earth Mover’s Distance (EMD): EMD^57^ measures structural differences between two distributions. Intuitively, earth mover refers to the amount of work needed to change one distribution to another, as if they are two piles of dirt. Here, we treat PCDs (of the original space data and DR embedding space data, respectively) as empirical one-dimensional distributions: in practice, we flatten the *N* × *C* matrices into vectors to represent observations from distributions that encapsulate the underlying global structure. Following the procedure of a previous usage of EMD as a global metric^20^, we perform min-max normalization of each vector to account for the difference in scale (original space PCDs can be on a different scale than embedding space PCDs). Subsequently, EMD is calculated after normalization. Thus using this metric, we are measuring the overall distributional differences between the original space PCDs and DR embedding space PCDs. In contrast with Spearman’s correlation, which captures the granularity of each cell’s positional change in terms of ranks, EMD treats the PCD distributions as a whole, providing insights on whether the embedding resembles the original data overall. To compute this metric, we used the SciPy implementation of the Wasserstein Distance with default parameters, which computes the distances between empirical cumulative distribution functions of the two vectors.

EMD and COR metrics are complementary to each other in the global category. Specifically, COR utilizes each individual cell’s relationship with all the others after DR, which is one perspective of the Global structure. On the other hand, EMD offers a zoomed-out perspective by treating all the distances as a distribution rather than distinguishing among each one of them individually. In practice, we believe that both approaches are important and highly complementary with each other.

### Accuracy - local structure preservation

We focus on neighbors for local metrics. Namely, we aim to assess whether each cell’s neighbors in the original space are still neighbors after performing DR. This category is important for not only clustering and cell type assignment but also understanding relationships between cells. The conventional approach of finding neighbors using BallTree or KDTree in sklearn proves to be quite slow for large datasets in the case of CyTOF. To realistically benchmark numerous methods on a large number of samples while also offering a quick interface for end users, we chose an approximate but fast algorithm called Approximate Nearest Neighbor Oh Yeah (ANNOY, github.com/spotify/annoy). It utilizes a locality-sensitive hashing algorithm^58^ to approximate distances along with using trees to divide data into subspaces, which are represented by a random sample of points, for fast nearest neighbor computations. As with the case of PCD, ANNOY offers a good, practical compromise for CyTOF data in terms of efficiency and accuracy. In our benchmarks, we found that the number of neighbors, *k* = *100*, works well in general: it accounts for the large sample size of CyTOF without adding a significant amount of computation time.

*K*-Nearest Neighbors (KNN): For each cell in the original and embedding space, we found *k* nearest neighbors using ANNOY, excluding the cell itself. We define neighborhood preservation by calculating the percentage of neighbors that overlap in both sets. Specifically, for each cell, we took the intersection of the two neighborhood sets from the original and embedding space, and the percentage of preserved neighborhood was calculated by dividing the cardinality of the intersection set by *k*. The resulting percentage was subsequently averaged across all cells. A larger KNN score close to 1 indicates nearly perfect alignment of neighbors, whereas a relatively small score shows poor neighborhood preservation. Importantly, here we are not using the machine learning algorithm of KNN to define cell types or cell clusters, as in the Random Forest Cluster Prediction metric below. So it is not to be confused with the RF metric below, which is a downstream analysis performance metric.

Neighborhood Proportion Error (NPE): As proposed by Konstorum *et al*^59^, NPE measures how many of the neighboring cells of the same class are preserved. This metric takes into consideration the classification of cells in neighborhoods, which are useful indications of whether clusters are well-separated after embedding. We implemented NPE in house using the following algorithm:

1. We defined *k* = *100* neighbors for each cell in the original and embedding space.
2. For each cell, we found the number of neighboring cells that belong to the same cluster. We used the original space clustering in this step, and again assume that there are *C* clusters. This results in two vectors of length *N*, one for the embedding space and one for the original space, where each element is the count of neighboring, same-cluster cells.
3. The two count vectors in Step 2 were divided by *k* = *100* to obtain the proportion normalized with the neighborhood size.
4. For cells belonging to each cluster *c*, we performed the following:

a. We subsetted in the normalized count vectors with the corresponding cells. This yields subsetted vectors of length *N*_e_, where *N*_e_ is the number of cells in the given cluster.
b. Using similar notations with the original authors, we found the kernel density estimates of the vectors from Step 4a, which we call **P**_**c**_ and **Q**_**c**_respectively, where subscript *c* indexes cluster. In the rare cases that **P**_**c**_ and **Q**_**c**_are inestimable point mass, this cluster is skipped, and we continued to Step 4a for the next cluster.
c. Finally, we calculated the the Total Variation Distance between the distribution for each cluster using the formula *sup*_a∈[_*_0,1_*_]_ |*P*_c_(*α*) − *Q_c_*(*α*)|.
5. The total variational distances were then averaged across all clusters to produce the NPE metric.

As a distance-based metric, smaller NPE indicates less error, meaning that each cell’s neighboring cells have similar composition in the original and embedding space. This method belongs to Local instead of Downstream, because it is focused on neighborhoods of individual cells instead of clusterings and cell types on a larger scale.

KNN focuses on whether the exact cells in the neighborhood of each cell are preserved between the original data and the DR space, while NPE focuses on whether the neighborhood of each cell in the DR space consists of cells of the same type as this cell, with the “type” or “cluster” of cells defined in the original data. Roughly speaking, KNN operates at the cell level, while NPE works at the cluster/cell type level. They are very complementary with each other overall.

### Accuracy - downstream analysis performance

A common first step of the downstream analysis pipeline is clustering and subsequently cell type assignment. The Downstream category specifically examines how each DR method affects the clustering performance, which will in turn affect other downstream workflows. Downstream has three sub-categories: Cluster Reconstruction measures how well original space clustering performs on the embedding space data; Cluster Concordance examines clusterings of the original space and the embedding space at the same time; and Cell Type-Clustering Concordance compares manually assigned cell types and embedding-space clustering. For the latter two sub- categories, the same metrics are used, but their purposes, inputs, and interpretations differ.

Cluster Reconstruction: Custer Reconstruction consists of four metrics to assess how the original space clustering performs after DR. While a clustering algorithm optimizes for the original data, the results should be reasonable if a DR method faithfully captures the structure of the original. This is often how such clusterings are visualized. In this sub-category, there are four equally weighted metrics: the first three measure the clustering quality itself whereas the last utilizes the clustering as a practical metric.

Silhouette Score^15^ is a measure of how cells are similar, in terms of Euclidean distance, to other cells in the same cluster as compared with the cells in other clusters. A large score close to 1 indicates a good clustering with tight clusters, where on the other hand a negative score means that the cells in clusters are more similar to other cells than those in the same cluster. Davies-Bouldin Index^60^ measures the average maximum ratio of intra-cluster distance over inter-cluster distance. A good clustering with a good embedding should result in clusters that are both tightly- formed and distinctly far apart, thus resulting in a small score. Calinski-Harabasz Index (CHI) measures variance instead of distance. Specifically, CHI is proportional to the ratio of inter-cluster variance as computed with cluster centroids and the centroid of the dataset over the intra-cluster variance computed with points in the cluster and each cluster centroid.

We also incorporated Random Forest Cluster Prediction (RF) to assess how well a random forest classifier can predict the cluster labels using embedding space data. As proposed by Becht *et al*^27^, the prediction accuracy is a practical metric to assess whether original clusters are distinguishable. We split the embedding data into 67% training data and 33% testing data. For each DR method, we trained a random forest classifier using sklearn and recorded the prediction accuracy using the testing set. Obviously, a higher accuracy is desired.

Cluster Concordance: Instead of focusing on original space clustering, the Cluster Concordance sub-category uses Adjusted Rand Index (ARI)^61^ and Normalized Mutual Information (NMI)^62^ to measure how clusterings of the original and DR space align. Given two clustering labels, a pair confusion matrix can be formed by considering every pair of data. The original Rand Index simply finds the ratio of correctly aligned results - the sum of true positive and true negative - over the total number of observations. The ARI adjusts the Rand Index with expected values. Intuitively, a higher ARI close to 1 means that the two clustering results are well aligned, suggesting that DR has lost little information. NMI, on the other hand, treats two clustering results as distributions, and the mutual information between them is computed. The resulting mutual information is then normalized by the entropy of the marginal. For NMI, a larger value close to 1 is preferred, whereas NMI of 0 means that the two clustering results are independent.

Cell Type - Clustering Concordance: While Cluster Concordance measures the efficacy of clustering after DR, a valid concern is whether such clustering is concordant with true cell types instead of arbitrary cluster labels. In this sub-category, we utilized cell types provided by the original authors when available as an independent source of true labels. If not, we manually assigned cell types using original space clustering. For good embeddings, we expect that cells of the same type are also clustered together, which also suggests that if practitioners use embedding space clusterings for cell type assignment, they will likely obtain reasonable results. We applied ARI and NMI to cell types and embedding space clusterings for each method. The algorithms are the same as the ones described in Cluster Concordance but with different inputs.

### Accuracy - scRNA-seq concordance

For datasets with matched CyTOF and scRNA-seq samples, analyzing these samples in conjunction is a great asset. One critical task in both workflows is DR, and as one would expect, DR results from matched samples need to reasonably be concordant. This will allow practitioners to not only employ a unified workflow for both but also make conclusions based on the same methods and assumptions. Further, many DR methods have indeed been used indiscriminately across different technologies. In this major category with three equally weighted sub-categories, we assessed the versatility of DR methods across technologies.

For this category, we assigned cell types for CyTOF samples using embedding space data produced by each DR method. scRNA-seq cell types were assigned with original space data after preprocessing. The goal is to assess which DR method’s results are the most concordant with the reference scRNA-seq data of the same sample.

Cluster Distance: This metric measures how the relationships between common cell types change between CyTOF and scRNA-seq samples. For example, if there are two closely related cell types with similar expressions, we expect them to be close to each other in both settings. To operationalize this concept, we first computed the centroid of each cell type for both CyTOF embeddings and scRNA-seq data, and only common cell types between two datasets are retained. Using permutation, we found all pairs of cell-type centroids. Let an arbitrary pair be cell type *a* and cell type *b.* Then, pairwise distance between *a* and all other centroids was computed and ranked. The rank of the distance between *a* and *b* was then normalized by the number of cell types and recorded. A small rank means that cell type *b* is close to the cell type *a* from the vantage point of *a*. The process is repeated for all pairs and for both the CyTOF embedding of interest and the scRNA-seq reference. One thing to note here is that the rank is not necessarily symmetric because, although the Euclidean distance between any *a* and *b* pair is symmetric, the rank obtained in this case depends on neighboring cell types. For example, If *a* has numerous neighboring cell types, whereas *b* is a lone outlier, the ranked distance between them from the vantage of *a* is large. But from *b*’s perspective, the rank can be much smaller. With each DR method, we have two resulting distance vectors of length *m*, where *m* is the number of permutations of cell-type pairs. Define **X** as the vector from a given CyTOF embedding and **Y** as the vector from the reference scRNA-seq sample. We then defined Cluster Distance as the L1 distance between **X** and **Y**.

Given that Cluster Distance is a distance-based metric, the interpretation is simple. A smaller value means that the relationships between cell types are well preserved. Specifically, relationship means the rank distance between cell types. The actual value of the Euclidean distance is not really meaningful given the different nature of CyTOF and scRNA data, but the rank captures the overall structure. This metric was implemented in house and available in CyTOF DR package.

EMD: Analogous to the contrast between Spearman’s correlation and EMD in Global, here the role of EMD is the same. In this metric, we use the same definitions of pairs of cell types as obtained from permutation. Instead of using only centroids, we calculate the distances between cell type *a* and all cells in cell type *b,* which are subsequently ranked and normalized by sample size. The same procedures are performed on both the CyTOF embedding of interest and the scRNAseq reference. This more fine-grained approach is possible due to the fact that EMD compares two distributions. Using the same notation as Cluster Distance, here we compute the EMD between **X** and **Y**.

We prefer a smaller EMD to suggest good cell-type relationship preservation on a global scale. In this case, the distribution of ranks is being measured **X** and **Y**. As with EMD in Global, this metric offers a good sense of whether the overall structures of cell types as clusters are well preserved using a DR method in CyTOF.

Gating Concordance: Gating Concordance is a validation metric to ensure that the cell type assignment in CyTOF makes sense for each DR method. Specifically, we manually assigned cell types using both original and embedding space data. For the latter, cell types were assigned using clusterings based on embeddings of each DR method. As with other concordance categories, we used ARI and NMI to assess whether embedding space cell types align well with original space cell types. This sub-category is warranted in the scRNA-seq Concordance category because the quality of embedding space cell type assignment is critical. If Gating Concordance is poor, then the results of EMD and Cluster Distance are not as meaningful by assumption. Thus, for a method to have good scRNA-seq Concordance, we expect it to excel in all three categories.

### Stability

We define stability as whether a certain method’s embeddings remain similar while executed on similar datasets. We bootstrapped Sample A of the Oetjen cohort 100 times and then executed all DR methods on the 100 bootstrapped samples (N=147,570). For the reference embedding from the original dataset and all the embeddings from bootstrapped datasets, we computed the PCD for each. As previously described, the PCD algorithm results in a *N* × *C* distance matrix, where *K* is the number of clusters or cell types. For the stability measure, we treated each PCD matrix as a single distribution by flattening it to become a vector of length *NC*. Without loss of generality, we denote these vectors as *P* and *Q*. To account for the scale difference across different DR methods, we performed min-max normalization to **P** and **Q**. We then used the EMD to measure the difference in distribution to quantify the structural difference *via* difference in distribution, which can be written as *EMD*(*minmax*(**P**), (*minmax*)(**Q**)). In essence, **P** is the reference embedding for each method, whereas each *Q* from the bootstrapped iterations is compared against the reference **P**. For each DR method, the computed distance from the 100 iterations were averaged to form an overall stability measure. Smaller EMD indicates that a given method’s embedding is structurally similar to the original embedding from run to run, and vice versa. DiffMap reported errors on all runs of this benchmark, likely due to the resampling scheme. Spectral was unable to run on the original dataset.

### Scalability

We measured each method’s runtime and memory consumption to assess scalability in a two-faceted experiment. To best assess performance on real datasets in a controlled manner, we used Sample B and H of the Oetjen cohort with 34 protein markers as the basis. Then, we employed random up- and down-sampling to comprehensively simulate both the number of cells and the number of different protein markers. By fixing the number of markers at 34, we subsampled without replacement 2,000, 5,000, 8,000, 10,000, 20,000, 30,000, 40,000, 50,000, 100,000, 150,000, 200,000, 300,000, 500,000, 750,000, and 1,000,000 cells. For the number of protein markers, we sampled 10, 20, 30, 40, 50, and 60 features, while fixing the number of randomly sampled cells at 100,000. In the latter case where up-sampling is needed, we utilized the original 34 markers as is and then randomly sampled the remaining markers from these 34 markers with some added random noise. No further method-based subsampling was allowed in this benchmark because doing so would defeat the purpose of assessing scalability fairly across all methods. For methods that would abort at sample size of 100,000, which is a reasonable sample size from a typical CyTOF cohort, we could only report error in the case of protein marker benchmark (**Fig. 5f**).

We performed all scalability benchmarks on the server with the same hardware configuration as detailed in the “Benchmark environment” section. Runtime and memory usage were measured on the Operating System level with the “/usr/bin/time -v” command in Linux. We used the “Elapsed (wall clock) time” as the measurement of runtime and “Maximum resident set size” as memory usage. Each measurement consists of the benchmark for one method on one dataset configuration, and all runs were sequential to allow methods to utilize as much system resource as possible without aborting. All other variables, such as file input and output, were kept the same across all runs and their impact was overall negligible. We then calculated and reported runtime in minutes and memory in GB for convenience.

### Usability

We defined usability with 13 criteria in five categories related to code quality and user experience. For each method, we manually inspected the quality of documentation, level of software support, the interface available to users, and implementation details that would benefit practitioners. All information is gathered from their websites, GitHub repositories, in-software docstrings and examples, and our own benchmark processes. We scored all methods using a binary scoring system: if a criterion is met, the method receives a score of 1; otherwise, it receives 0. Under circumstances where ambiguity may occur, such as whether the package has been updated as needed, we generally give such edge cases the benefit of the doubt unless clear incompatibility or lack of support is present. The final ranking of usability is based on the sum of scores received by each method (**Fig. 6b**). The exact usability criteria are detailed below.

Documentation: (1) Examples: The project repository or website provides examples for users to follow. (2) Website: The method has a dedicated website for detailed documentation.

Implementation: (1) CLI: The method has a command-line interface that is capable of directly running the DR. (2) Cross platform: The method is easily performed in both R and Python.

Installation of packages and alternative implementations are allowed. (3) Import: The package can be imported within its respective software environment. This allows users to integrate methods into their existing workflow without resorting to custom command-line scripts. (4) Matrix: The method takes a general expression matrix or array as input instead of requiring a specific data structure.

Maintenance: (1) CI/CD: The package has an automated workflow to test codes with new development. (2) Update: The package has been updated within the past year or as needed.

Publication: (1) Built-in: The implementation of the method is built into R or Python without needing any installation. (2) Open Source: The package is open source. (3) Published Package: The package has been published on a standard package repository, such as PyPI, CRAN, or conda.

Usage: (1) Mapping: The method has a mapping function that can map new data onto existing embeddings. (2) Robust: The method does not produce unexpected exceptions or errors, except for sample size limitations.

### Real CyTOF datasets used in this study

We accessed a total of 11 cohorts of 110 real CyTOF samples for this benchmark study (**Supplementary Data 2**). These include the TuPro cohort^10^, the Oetjen cohort^33^, the CyAnno cohort^63^, the LG cohort^64^, the Brain cohort^65^, the BC cohort^66^, the Levine32 cohort^67^, the Levine13 cohort^67^, the Samusik cohort^68^, the Lung cancer cohort^69^, and the Covid cohort (CyTOF_00000000000001 to CyTOF_00000000000012 on DBAI). For all datasets, we either obtained their cell typing results from the original publications, or performed cell typing ourselves based on clustering of the cells followed by manual assignment. We showed the manual cell typing results of the Lung cancer cohort, in **Supplementary Fig. 5**, as an example. Matched scRNA-seq data are available for the TuPro and Oetjen datasets. For the BC dataset, there were too many samples. To avoid biasing our benchmarking results, we selected the top 14 datasets with more than 5,000 cells.

For all samples, we deployed our standardized preprocessing pipeline (included as part of CyTOF DR Package) unless they have been preprocessed beforehand by the original authors. Our pipeline includes identifying lineage channels, ArcSinh transformation with a co-factor of 5, gating to remove debris using instrument channels, and bead normalization. In general, we did not perform cross-batch normalization to avoid any bias introduced by such algorithms. For the cases where the data have been previously processed, such as the CyAnno cohort, we manually inspected the data and applied specific steps of the pipeline on an as-needed basis: if cell types have already been assigned, we did not gate to remove debris; when not all bead channels were available, we assumed that bead normalization had already been performed on instrument. For all samples, we cross-checked the preprocessing results with DR embeddings of the cells labeled with their cell types.

### CyTOF data simulation (*Cytomulate*)

We devised a probabilistic simulation model that captured the key features of the arcsinh(./*5*)- transformed CyTOF data. Suppose that the desired resulting dataset contains *m* = *1*, ⋯ , *M* protein markers, *n* = *1*, ⋯ , *N* cell events, and *p* = *1*, ⋯ , *P* cell types. Let the probability of observing the *p*th cell type be *π*_p_. Then, for the *n*th cell event, we can construct a cell type indicator variable *Π*_n_ ∼ Categorical(**π**), where **π** = [*π_1_*, ⋯ , *π*_*P*_] is a *P*-by-1 probability vector.

Given the cell type *p* for the *n*th cell event, we assume that, without cell differentiation or background noise, the expression level for the *m*th marker is 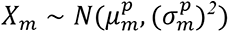. Since a cell type typically expresses a set of marker genes, to further mimic the characteristics of CyTOF data, we associate, with each cell type *p*, a set *A*^ϕ,p^ = {*m*: the *m*th marker is not expressed in cell type *p*}. Then, given *A*^ϕ,p^, we set 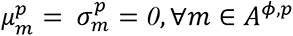.

To simulate cellular lineage, we assume that the underlying cell lineage is composed of several non-overlapping cell differentiation trees, where cell type *p* differentiates into *z*^p^“children” *p*^e^^1^ , ⋯ , *p*^e^z^p^. From the *m*th marker of each cell type *p* to the *m*th marker of each of its “children” *p*^e^, we construct a differentiation path 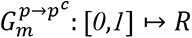, such that 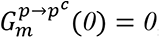, and 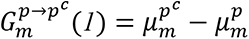, where 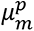 and 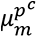 are the mean expression levels of the *m*th marker of cell type *p* and cell type *p*^e^, respectively. We also assume that 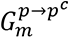 is a stochastic process, whose realization is denoted as 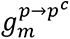.

With these setups, the simulated expression level for the *n*th cell event is calculated following these steps: (1) Generate a cell type *p* from Categorical(**π**). (2) If the cell type *p* has “children”, we randomly select from them a cell type *p*^e^. (3) For each marker *m*, we generate a sample *x*_m_ from 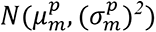, a “pseudotime” 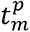 from Beta(*0*.*4*,*1*), and a background noise *e* from *N*(*0*, *σ*^2^). (4) The final expression level for the *m*th marker is then calculated as 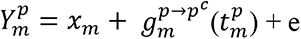.

### Subsampling for selected DR methods

In all accuracy benchmarks, we employed two strategies when a method fails due to large sample size. 1) If such a method has a mapping function to accommodate new data (e.g. LLE and Isomap), we initialized and trained the method with 10% subsampled data and then mapped the rest onto the embedding; 2) If a method has no mapping function and aborts during benchmarking, we treated such methods as “Error”, and they will be ranked last. This approach ensures that all evaluations are based on full samples rather than a mix of sample sizes. In a separate benchmark, we also executed all methods on subsampled data: if a method fails in this case, no further subsampling was allowed and they were treated as “Error”. For all subsampling in these benchmarks, we randomly sampled 10% of cells from each sample without replacement.

Although scvis does not have a memory constraint like LLE and Isomap, its authors recommend training the model using a small sample, as a unique feature of scvis. Thus, we performed scvis both with (denoted as *scvis*) and without subsampling (denoted as *scvis (Full)*). With the help of its mapping function, all evaluations were performed with complete samples.

### Development of the CyTOF DR playground webserver

We developed an online web tool for displaying the results of benchmarking DR algorithms on CyTOF data. The technical stack includes HTML, CSS, JavaScript and DataTables. The visualization table to display the benchmarking results of the real and the simulation datasets in this web tool was created with the DataTables jQuery plugin. This table provides searching, sorting and pagination features and progressive bars to visualize the performances of the DR methods according to different metrics. There are two main features in this web tool. The first one is to search for DR results based on multiple input conditions (left column). And the other is to dynamically calculate the average of the performance scores for the user-chosen metrics and to rank the chosen DR methods based on the average scores (top row). This web tool is hosted on the Database for Actionable Immunology (DBAI) website (https://dbai.biohpc.swmed.edu/)47,48,70.

### Statistics & reproducibility

Computations were mainly performed in the R (3.6.3 and 4.1.3) and Python (3.7 and 3.8) programming languages. All statistical tests were two-sided unless otherwise described. We used the nonparametric Wilcoxon Signed Rank Test with “wilcox.test” and “pairwise.wilcox.test” procedures in R for testing ranking differences between samples in terms of accuracy. All pairwise tests’ p-values were adjusted with the Benjamini-Hochberg Procedure to control the false discovery rate (FDR). For correlation, we used Spearman’s rank-based correlation coefficient.

For all rank-based metrics and results, higher ranks indicate superior performance. In case of tied ranks, we employed the maximum rank to allow tied values to all have the highest rank among them. The highest possible rank is 21 while the lowest is 1, unless otherwise noted such as in the case of parameter tuning. Decimal ranks are results of the weighting scheme of our evaluation framework.

No statistical method was used to predetermine sample size. We used all CyTOF samples from the original sources, except for the Imaging CyTOF cohort. There were too many samples in this cohort. So to avoid biasing our benchmarking results, we selected the top 14 datasets with more than 5,000 cells. This study was not a clinical trial, so the experiments were not randomized. The investigators were not blinded to allocation during experiments and outcome assessment.

## DATA AVAILABILITY

The public CyTOF and scRNA-seq data used in this study were from citations^10, 12, 33, 64–69, 71–73^ and described in **Supplementary Data 2**, with more details on data accession documented in this table. The in-house COVID-19 vaccinee CyTOF datasets were available from DBAI (https://dbai.biohpc.swmed.edu/), with accession codes from CyTOF_00000000000001 to CyTOF_00000000000012 (https://dbai.biohpc.swmed.edu/cytof-database.php). Source data are provided with this paper.

## CODE AVAILABILITY

The CyTOF simulation model is available at: cytomulate.readthedocs.io. The CyTOF DR Package webserver is available at CytofDR.readthedocs.io. The CyTOF DR Playground webserver is available at dbai.biohpc.swmed.edu/cytof-dr-playground.

## ACKNOWLEDGEMENTS

None

## AUTHOR CONTRIBUTIONS STATEMENT

K.W. performed all DR benchmark analyses and created CyTOF DR Package. F.W. and K.W. created CyTOF DR Playground. Y.Y. and K.W. created *Cytomulate*. Y.Y. provided feedback on evaluation metric implementations. B.S. provided codes for pseudotime inference. X.W. conceived the study. T.W. and X.W. designed and supervised the whole study. All authors wrote the manuscript.

## COMPETING INTERESTS STATEMENT

Dr. Tao Wang is one of the scientific co-founders of NightStar Biotechnologies, Inc.

## TABLES

None

## FUNDING SOURCES

This study was supported by the National Institutes of Health (NIH) [5P30CA142543/TW, 1R01CA258584/TW, XW, U01AI156189/TW], and Cancer Prevention Research Institute of Texas [RP190208/TW, XW, RP230363/TW].

## Notes

### Competing Interest Statement

The authors have declared no competing interest.

### Summary of Updates

This revision consists of formatting changes and clarifications.

