## Supplementary Information for "Comparative Analysis of Dimension Reduction Methods for Cytometry by Time-of-Flight Data"

**Supplementary Fig. 1 Number of yearly citations returned by Google Scholar, by searching for "CyTOF", "scRNA-seq", and "CyTOF & scRNA-seq".** The blue curves show the true citations counts, and the orange curves show the exponential curves fitted to the true citation counts. Source data are provided as a Source Data file.

**Supplementary Fig. 2 One example breast cancer Imaging CyTOF dataset.** The per-cell staining of the Ecadhe (E-cadherin), Ki67, SMA (α-smooth muscle actin), and CD45 channels were shown in the spatial context. Source data are provided as a Source Data file.

**Supplementary Fig. 3 Performances of the DR methods on the simulation data.** (a) Performances of the DR methods on the simulation data for each specific category of the accuracy metrics. (b) Performances of the DR methods on the simulation and real data for each major category of the accuracy metrics (scores of the specific categories averaged for each major category). Source data are provided as a Source Data file.

**Supplementary Fig. 4 The runtime and memory usage performances of several tSNE algorithms**. This plot focuses on the scalability of tSNE with increasing numbers of cells. "tSNE" in the main figures refers to "FIt-SNE (openTSNE)" here. "tSNE (sklearn)" in the main figures refers to "BH tSNE (Sklearn)" here. Source data are provided as a Source Data file.

**Supplementary Fig. 5 Manual cell type assignment results of the CyTOF data from the Lung cancer cohort.** All cell types are visualized by a colored UMAP embedding. Source data are provided as a Source Data file.

**Supplementary Note 1** Additional analyses and discussions involving the CyTOF DR methods

**Supplementary Discussion 1** The theoretical efficiency and scalability of the top DR methods, with respect to the dimensionality of CyTOF datasets.

Sup. Fig. 1

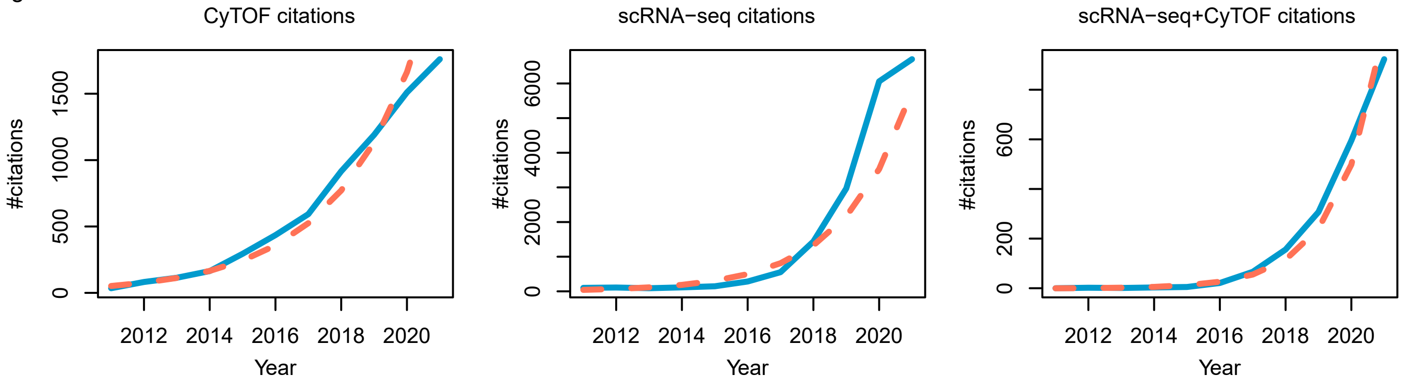

CyTOF citations
scRNA−seq citations
scRNA−seq+CyTOF citations
#citations
0
500
1000
1500
0
2000
4000
6000
0
200
600
2012
2014
2016
2018
2020
Year

Sup. Fig. 2

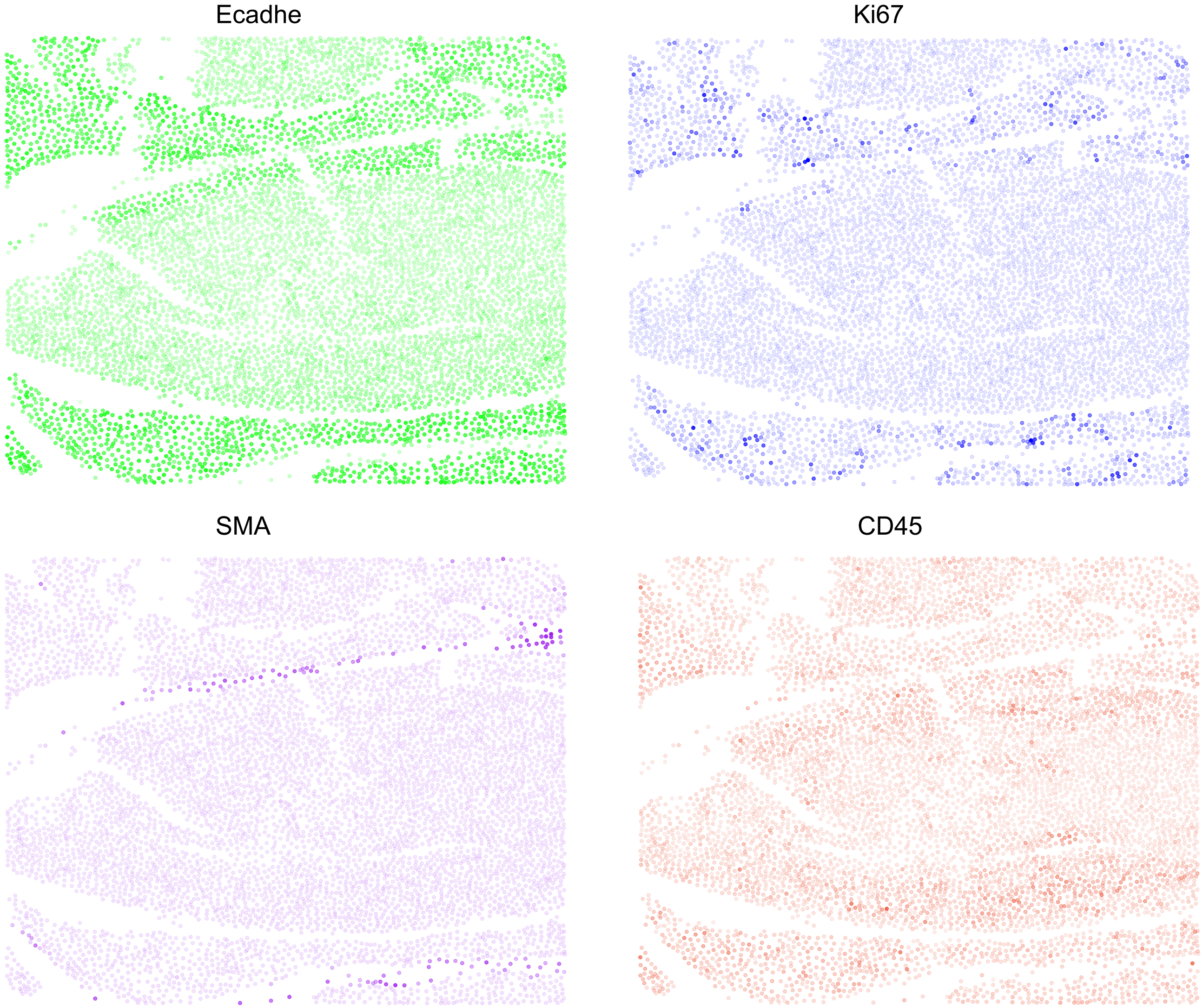

### Sup. Fig. 3

(a)

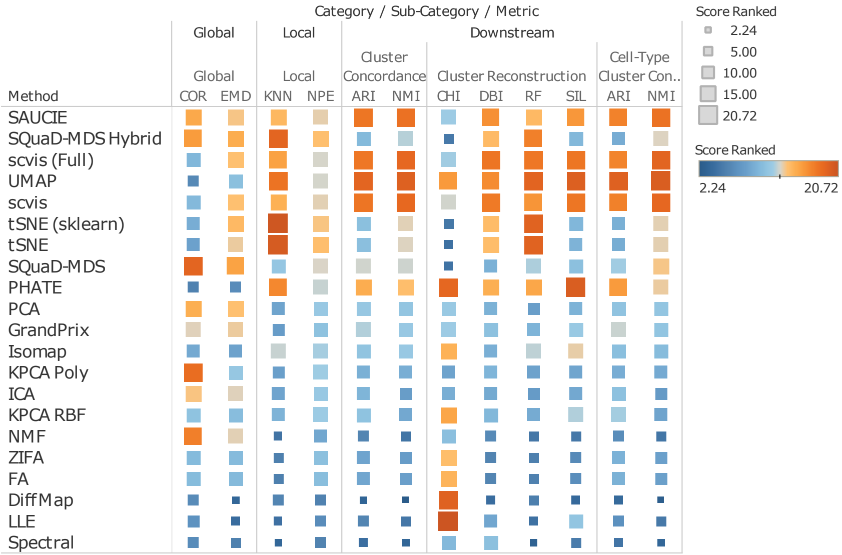

Category / Sub-Category / Metric
Global
Local
Downstream
Global
Local
Cluster Concordance
Cluster Reconstruction
Cell-Type Cluster Con..
Method
COR
EMD
KNN
NPE
ARI
NMI
CHI
DBI
RF
SIL
ARI
NMI
SAUCIE
SQuaD-MDS Hybrid
scvis (Full)
UMAP
scvis
tSNE (sklearn)
tSNE
SQuaD-MDS
PHATE
PCA
GrandPrix
Isomap
KPCA Poly
ICA
KPCA RBF
NMF
ZIFA
FA
DiffMap
LLE
Spectral
Score Ranked
2.24
5.00
10.00
15.00
20.72
Score Ranked
2.24
20.72


(b)

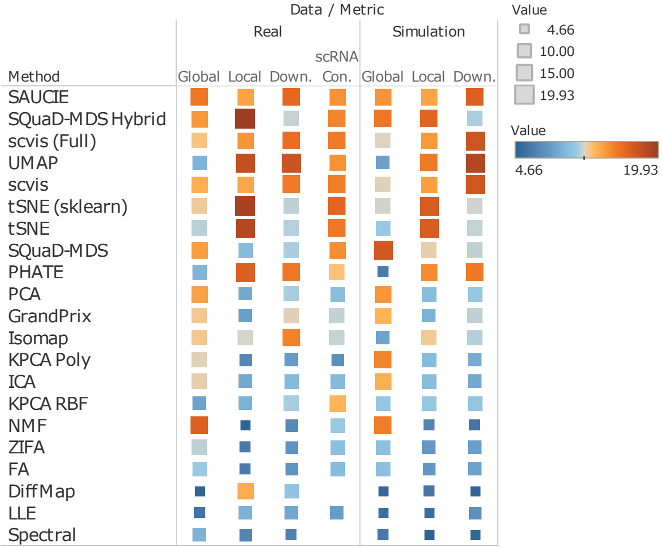

Data / Metric
Real
Simulation
Method
Global
Local
Down.
scRNA Con.
Global
Local
Down.
SAUCIE
SQuaD-MDS Hybrid
scvis (Full)
UMAP
scvis
tSNE (sklearn)
tSNE
SQuaD-MDS
PHATE
PCA
GrandPrix
Isomap
KPCA Poly
ICA
KPCA RBF
NMF
ZIFA
FA
DiffMap
LLE
Spectral
Value
4.66
10.00
15.00
19.93
Value
4.66
19.93

Sup. Fig. 4

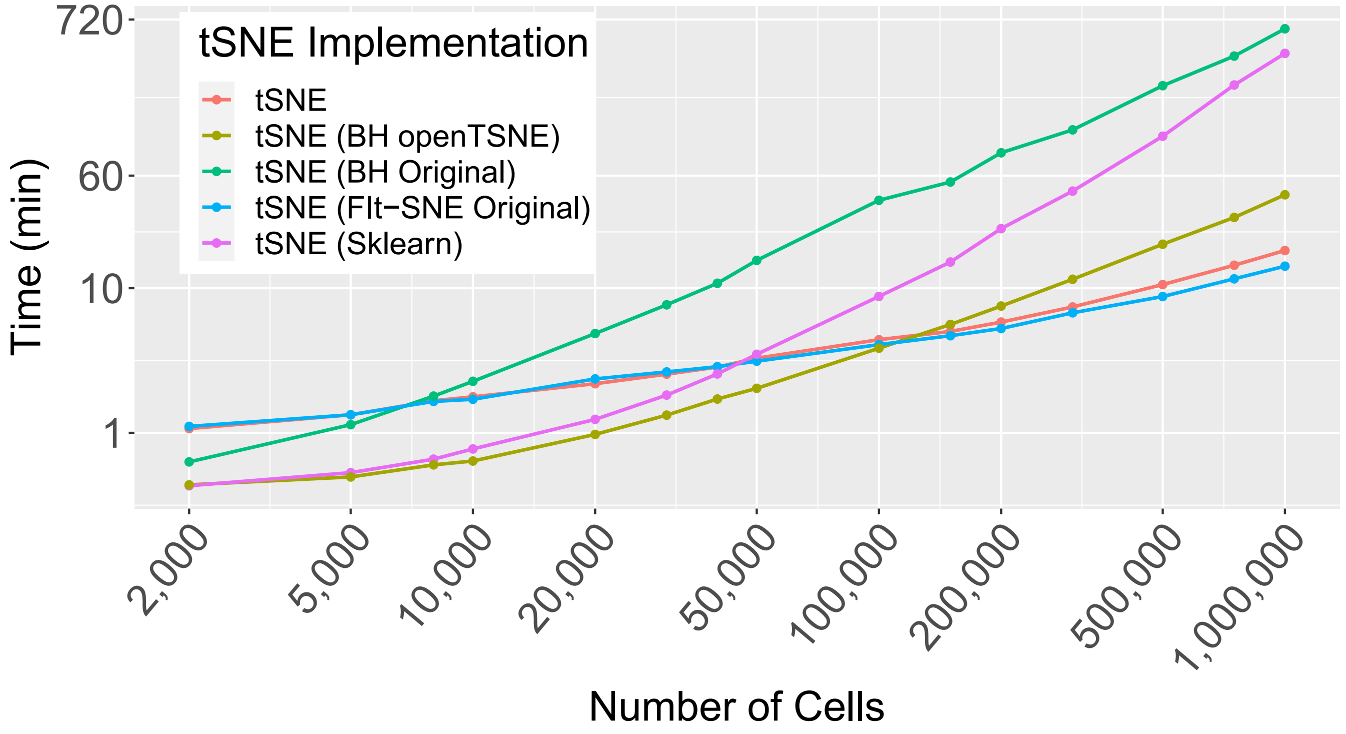

tSNE Implementation
tSNE
tSNE (BH openTSNE)
tSNE (BH Original)
tSNE (FIt−SNE Original)
tSNE (Sklearn)
Time (min)
720
60
10
1
2,000
5,000
10,000
20,000
50,000
100,000
200,000
500,000
1,000,000
Number of Cells


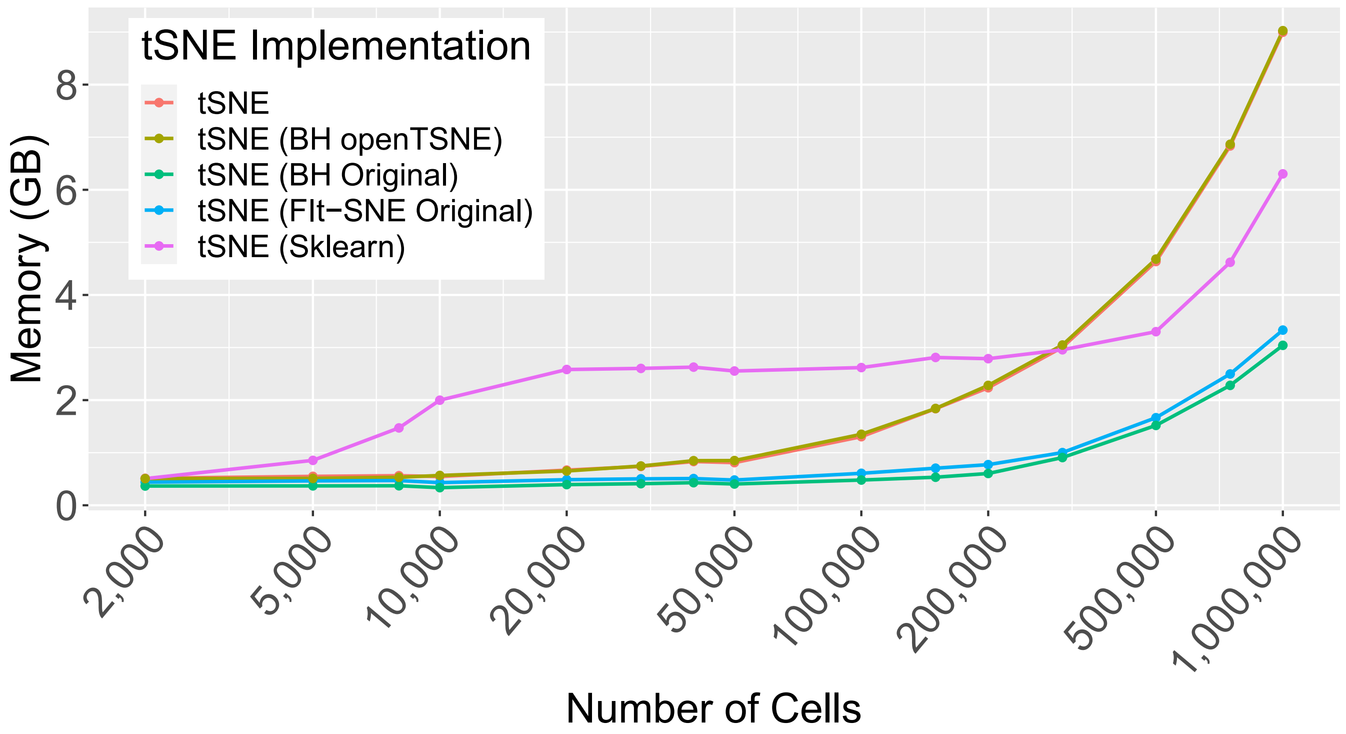

tSNE Implementation
tSNE
tSNE (BH openTSNE)
tSNE (BH Original)
tSNE (FIt−SNE Original)
tSNE (Sklearn)
Memory (GB)
8
6
4
2
0
2,000
5,000
10,000
20,000
50,000
100,000
200,000
500,000
1,000,000
Number of Cells

Sup. Fig. 5

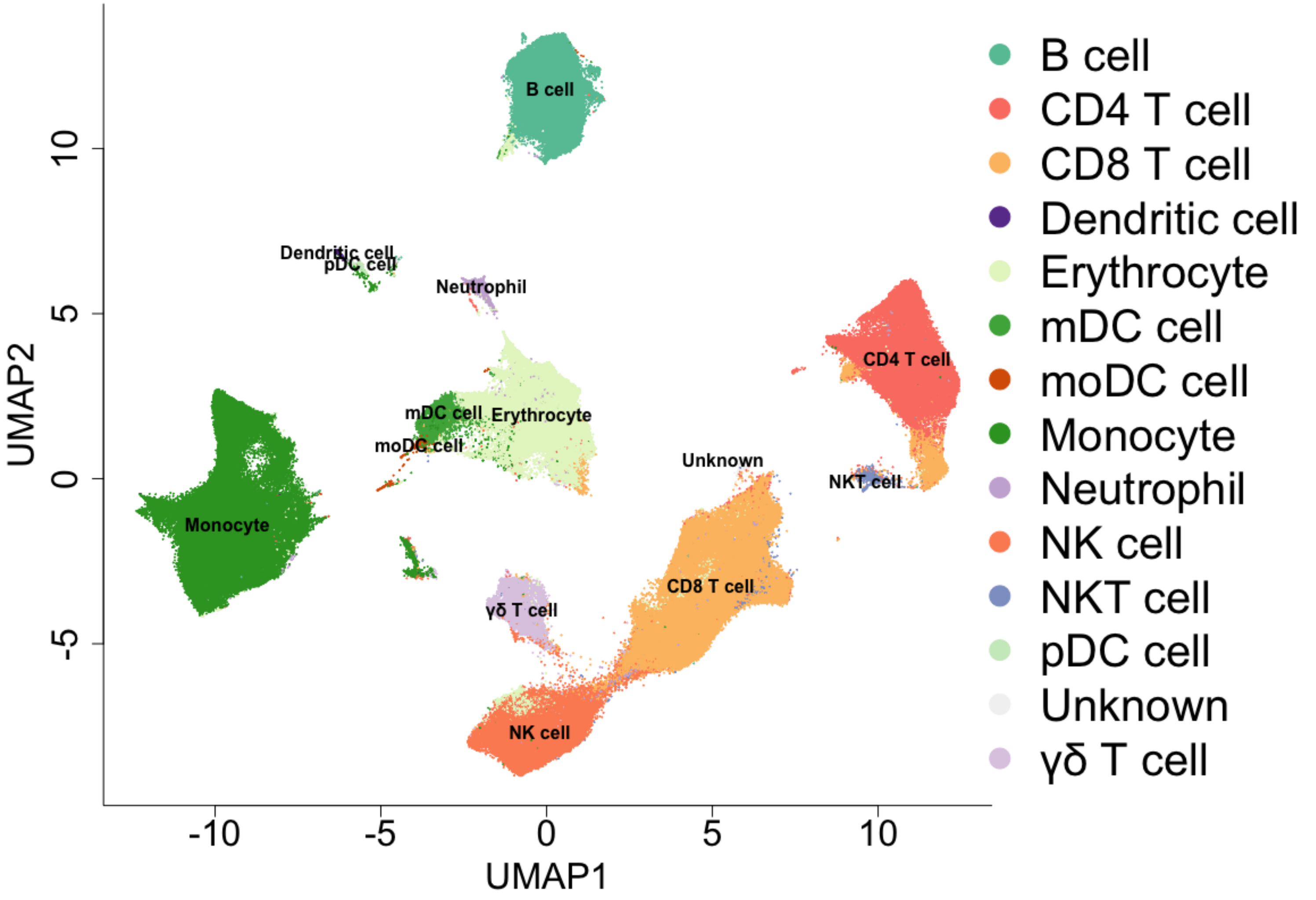
B cell
Dendritic cell
pDC cell
Neutrophil
CD4 T cell
mDC cell
Erythrocyte
moDC cell
Unknown
NKT cell
Monocyte
CD8 T cell
γδ T cell
NK cell
UMAP2
UMAP1
10
5
0
-5
-10
-5
0
5
10
B cell
CD4 T cell
CD8 T cell
Dendritic cell
Erythrocyte
mDC cell
moDC cell
Monocyte
Neutrophil
NK cell
NKT cell
pDC cell
Unknown
γδ T cell

### Supplementary Note 1: Additional analyses and discussions

#### Web resources we generated for this CyTOF DR review study

We generated three resources, a webserver for displaying the results for benchmarking CyTOF DR methods (**Supplementary Note 1 Fig. 1**), a webserver for providing our easy-to-deploy implementations of the DR methods and our evaluation metrics (**Supplementary Note 1 Fig. 2**), and the *Cytomulate* algorithm for simulating CyTOF data (**Supplementary Note 1 Fig. 3**).

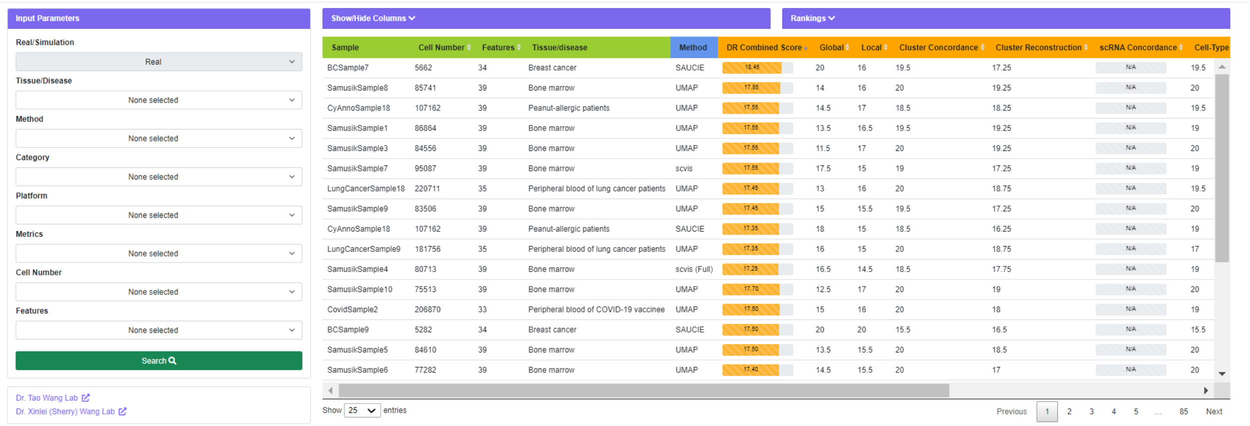


**Supplementary Note 1 Fig. 1 Screenshot of CyTOF DR playground.**

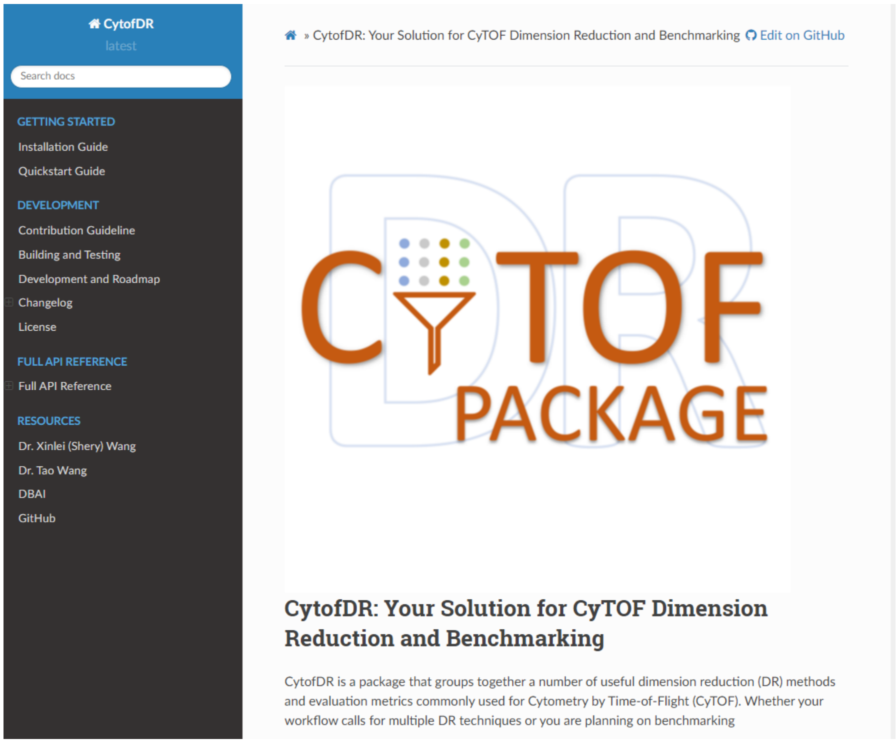


**Supplementary Note 1 Fig. 2 Screenshot of CyTOF DR package.**

To facilitate open and user-friendly deployment and evaluation of DR methods, we established the CyTOF DR Package website (CytofDR.readthedocs.io, **Supplementary Note 1 Fig. 2**) to provide implementation of the DR methods and our evaluation metrics, which is streamlined as much as possible. As we illustrated in **Fig. 1f**, a new DR user of CyTOF data can take a prior knowledge-driven approach of examining our DR benchmark results of the datasets evaluated in this study, shared conveniently through "CyTOF DR Playground", and then run their favorite DR methods using their own installation or the implementations that we provide on "CyTOF DR Package". The user can also take an empirical approach of deploying the evaluation metrics (or a subset of our evaluation metrics according to preference or data availability) on their own CyTOF datasets, using the evaluation and DR code implementation that we offered through "CyTOF DR Package", for picking the best approach. We also envision our system to be quite useful to future developers of DR methods for CyTOF data or other types of data in general. The extensive collection of evaluation metrics will allow future developers to craft their own benchmark recipes for DR in any field. At the same time, developers can easily extend the framework and add functionalities by simply deriving from our Python classes. We hope such a unified and streamlined evaluation-execution paradigm will become more widely adopted for various bioinformatics development scenarios.

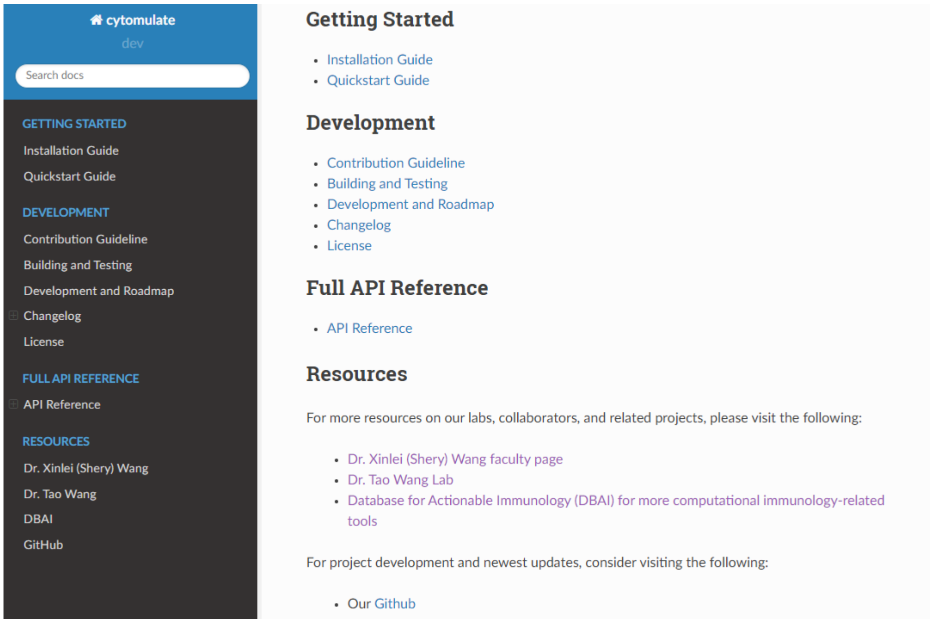


**Supplementary Note 1 Fig. 3 Screenshot of Cytomulate**

**An example simulated CyTOF dataset generated by *Cytomulate***

In **Supplementary Note 1 Fig. 4**, we showcase a simulated CyTOF dataset generated by *Cytomulate*. Multivariate Gaussians were used to generate the protein expressions of cells of various cell types. SAUCIE was used to visualize the single cells in a two dimensional space. According to the overall pattern of distribution of this dataset and those of real CyTOF data (**Fig.**

**4a**), we can see that *Cytomulate* closely mimics real CyTOF data. Compare the pattern of distribution of cells (such as level of dispersion with respect to the clusters) in this figure with the SAUCIE figure of **Fig. 4a**. More analyses and results validating the performance of *Cytomulate* in simulation of CyTOF data can be found in our *Cytomulate* work[1].

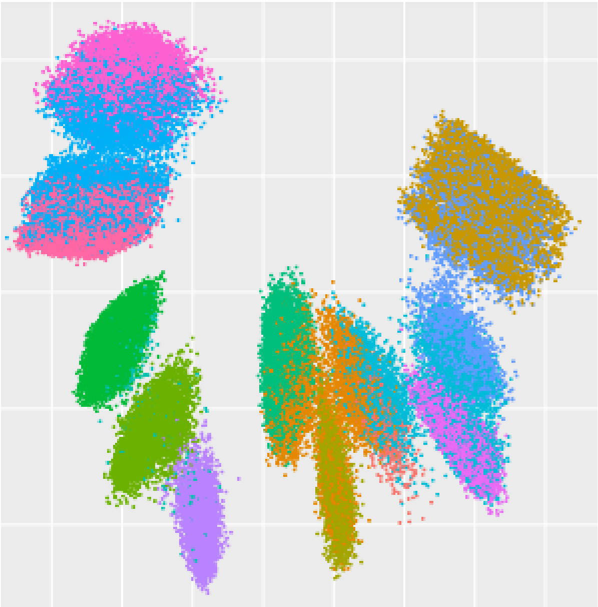

**Supplementary Note 1 Fig. 4 One simulated CyTOF dataset visualized in the SAUCIE space.** Source data are provided as a Source Data file.

**Detailed inspection of the DR results for the BC dataset**

SAUCIE is the top performer in terms of Global structure preservation. Further, SAUCIE significantly outperforms UMAP and scvis in the cell-type cluster concordance subcategory of Downstream performance (**Supplementary Note 1 Fig. 5a**). These two results suggest that SAUCIE is the best at detecting and distinguishing the main types of cells that exist in the samples. We classified the cells in the BC CyTOF samples into four clusters in the DR space as guided by the original BC publication, which reported four cell types (tumor, immune, vessel, stroma). Then, we visualized the clusters in the spatial context (**Supplementary Note 1 Fig. 5b**). The red cells in both panels and the green cells in the right panel from UMAP denote tumor cells. This shows that SAUCIE tends to be better at clustering all tumor cells together as a whole.

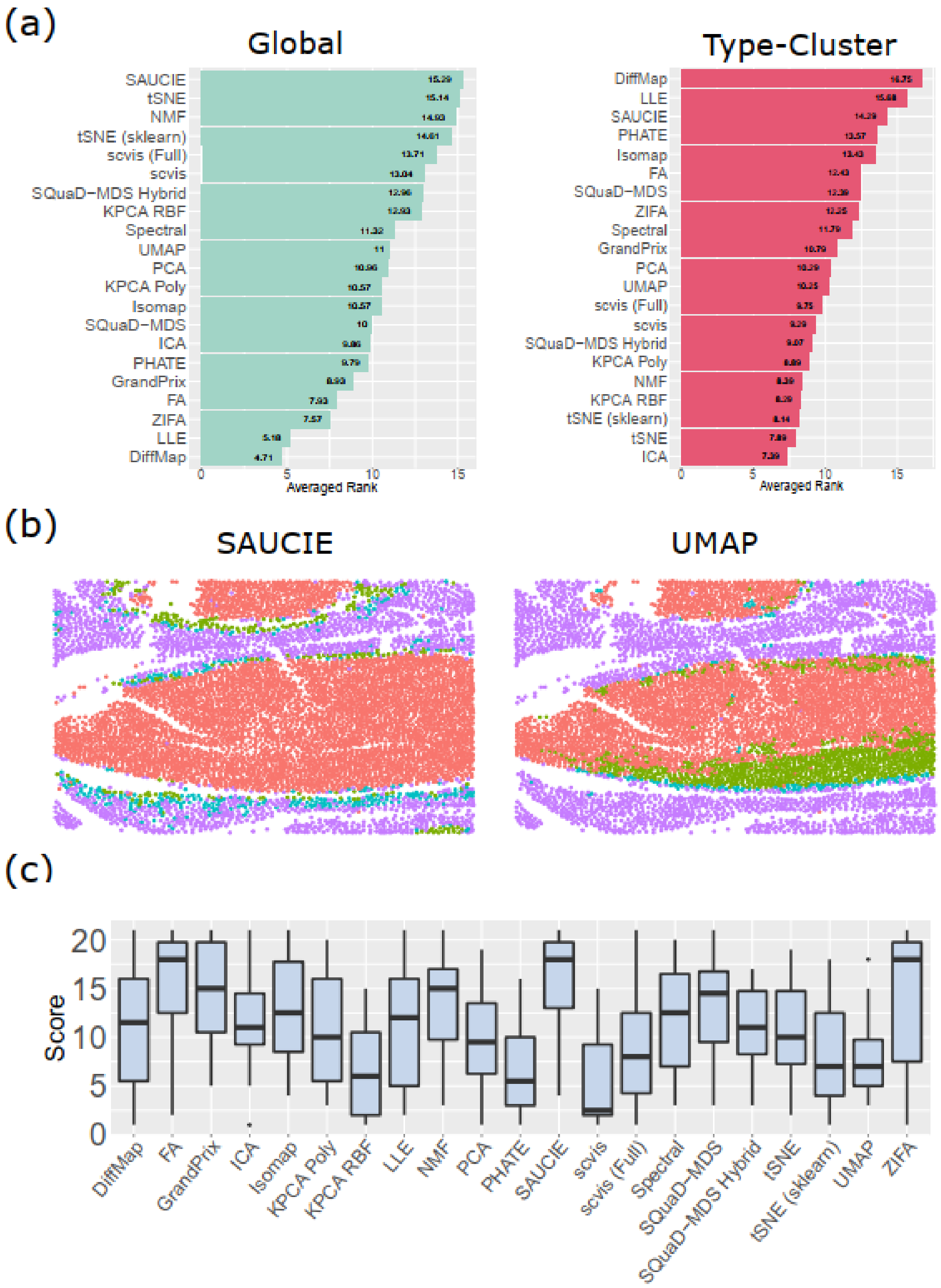


**Supplementary Note 1 Fig. 5 Closer examination of the DR results for the BC dataset.** (a) The performances of the DR methods in terms of global accuracy metric and cell type-cluster concordance subcategory of Downstream performance. (b) The cells in the BC dataset (sample #1) were classified into four types, according to clustering of the cells in the DR space. The cells and these clusters were visualized in the cells' spatial context. Left: SAUCIE; right: UMAP. (c) The correlation between spatial distances and the distances in gene expression between pairs of cells in the same "clusters" that were determined in (b). Box boundaries represent interquartile ranges, whiskers extend to the most extreme data point which is no more than 1.5 times the interquartile range, and the line in the middle of the box represents the median. For each method, the sample size is $N = 14$, which corresponds to the 14 samples from the BC cohort. Source data are provided as a Source Data file.

##### A comparison between downsampling-based and Point Cluster Distance (PCD)-based benchmark metrics

In our accuracy metrics, EMD and COR of the Global Category utilized PCD, which basically means that they used cluster centroids instead of actual data points and their embeddings to assess performance, in parts of the accuracy computation process. We chose this approach to avoid the quadratic computational complexity of certain metrics. However, one concern is that collapsing cell clusters to their centroid will obfuscate instances where two clusters overlap in the low dimensional embedding, and that clustering methods are not perfect, which means that a suboptimal clustering will lead to a suboptimal gold standard. To remove the concern regarding the negative impact of this potential caveat of PCD-based approaches, we also tested an alternative approach. We computed the pairwise distance in the embedding and DR space in a randomly sub-sampled set of 10,000 cells (without replacement) for each CyTOF dataset and then computed the full pairwise Euclidean distances within this subset. Other ranking and analysis methods remained the same, as they did not involve PCD. We compared the results obtained from both approaches in **Supplementary Note 1 Fig. 6** and found that both PCD- and subsampling-based methods yield very similar results in real and simulated CyTOF samples. Critically, the results are nearly identical for the top-ranking methods.

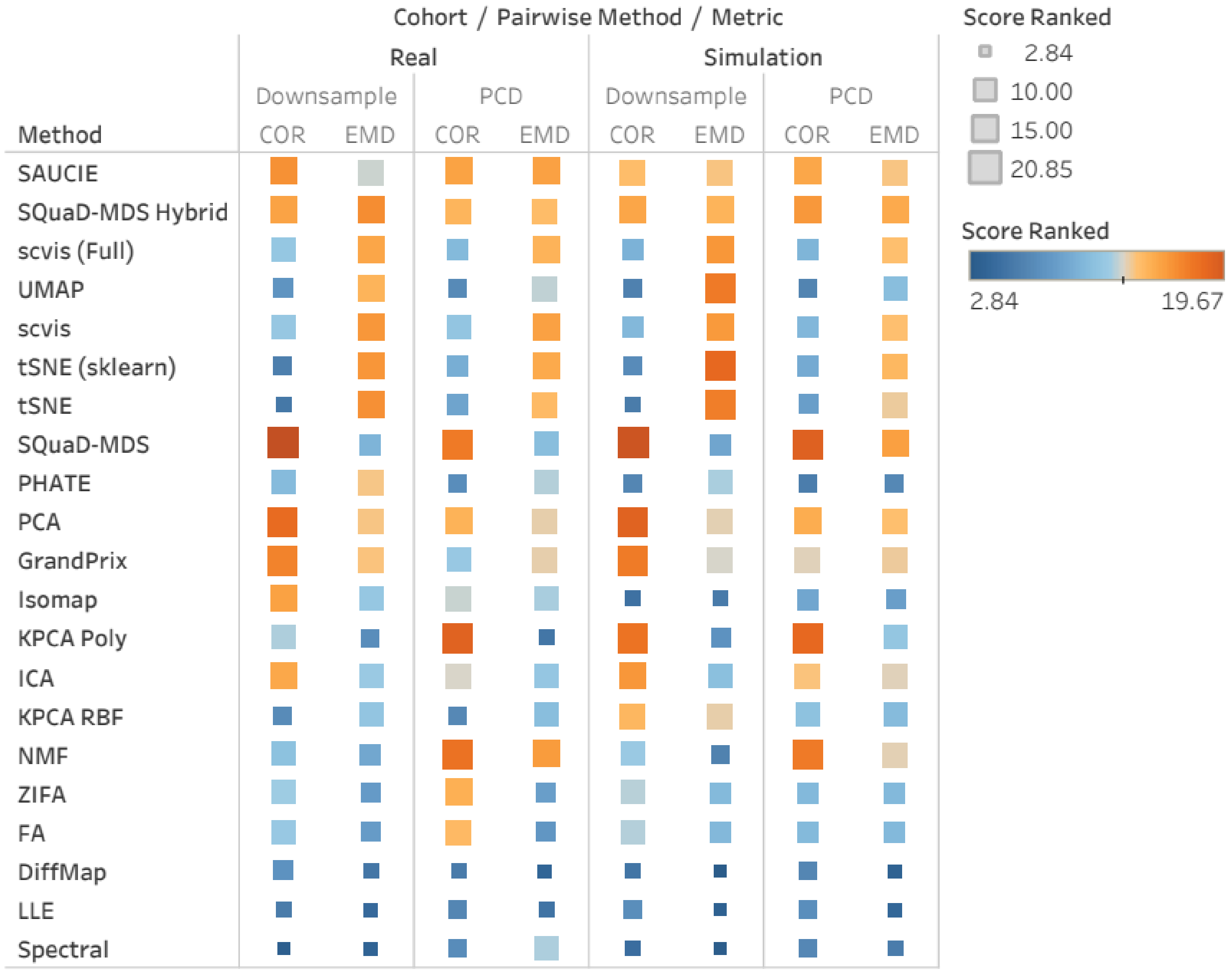


**Supplementary Note 1 Fig. 6 Global accuracy results from both downsampling-based and PCD-based approaches.** Source data are provided as a Source Data file.

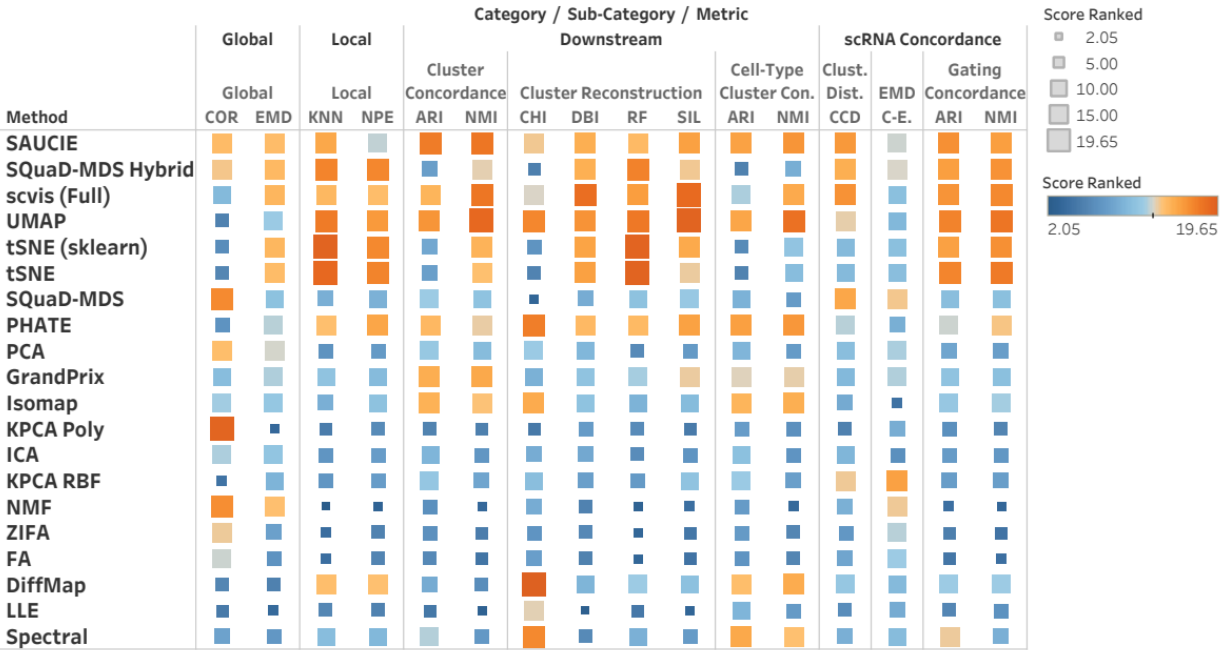


**Supplementary Note 1 Fig. 7 Accuracy performance on all 10% downsample real datasets.** This plot corresponds to the results displayed in **Fig. 3b**, which does not downsample for evaluation. The performance of DR methods is similar to that of the main figure. Source data are provided as a Source Data file.

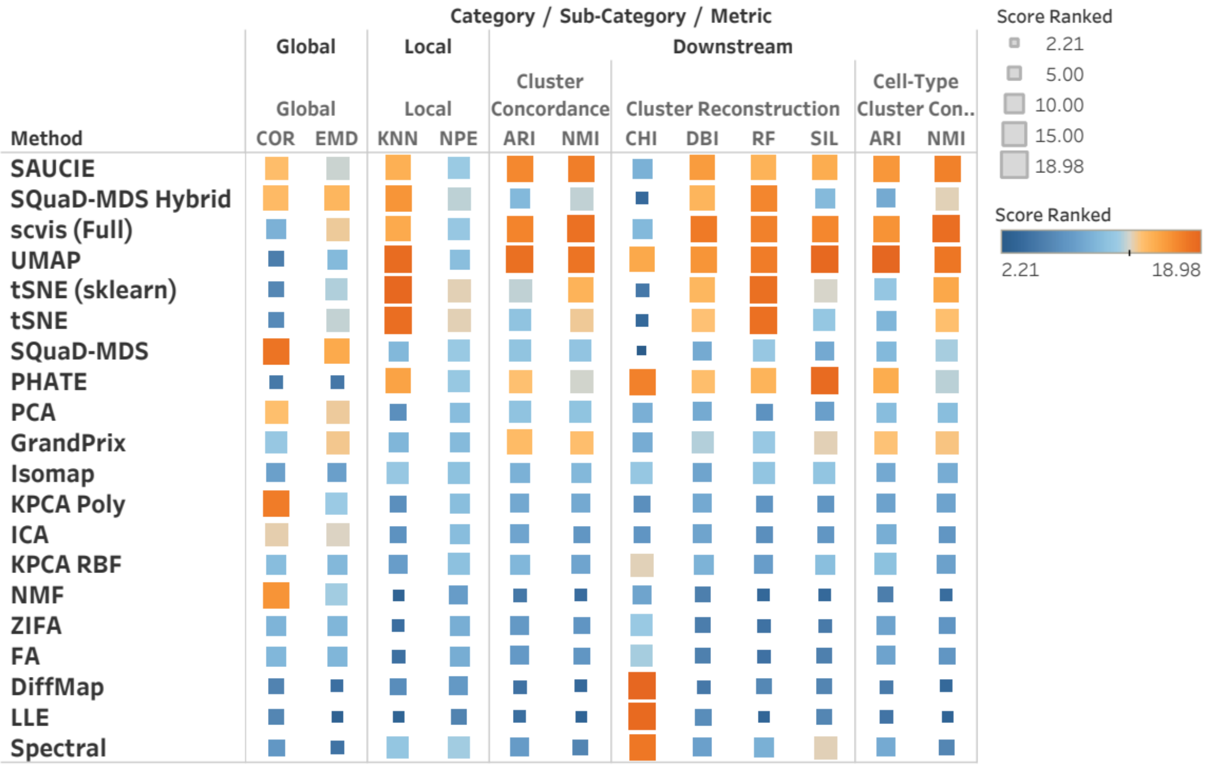

**Supplementary Note 1 Fig. 8 Accuracy performance on all 10% downsample simulation datasets**. This plot corresponds to the results displayed in **Supplementary Fig. 3a**, which does not downsample for evaluation. As with real datasets, the performance shown here is again comparable. Source data are provided as a Source Data file.

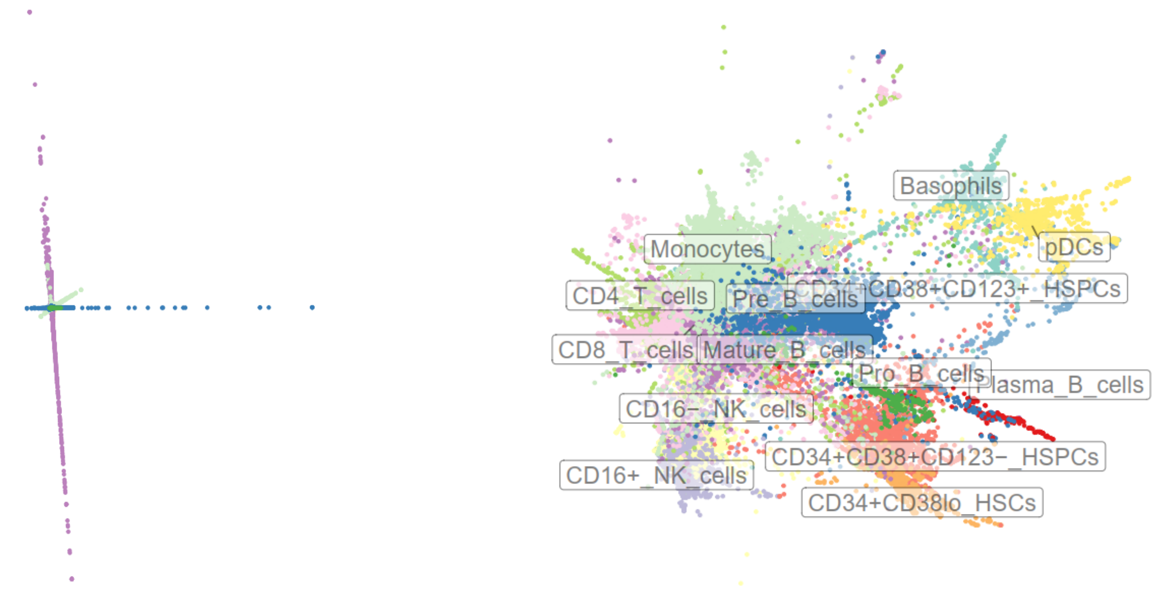


**Supplementary Note 1 Fig. 9 Examples of bad DR embeddings from the Levine32 dataset.** The left panel is LLE, which fails to distinguish any cell types or clusters meaningfully. The right panel is the PHATE embedding. PHATE, which usually performs decently, produces a confusing embedding with cell types mixed with each other. Source data are provided as a Source Data file.

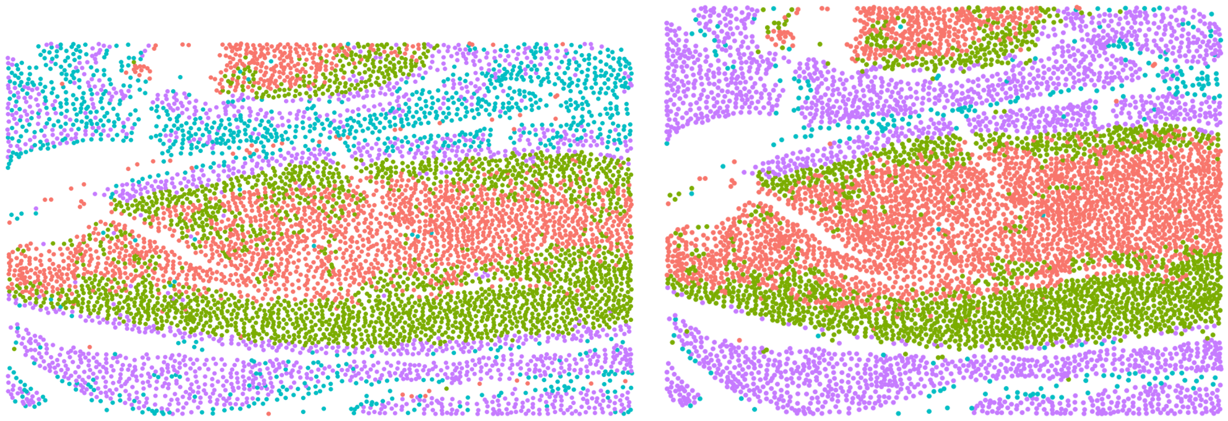

**Supplementary Note 1 Fig. 10 Examples of bad IMC plots from the BC cohort Sample 1.** The left panel is clustering results from KPCA RBF and the right panel from scvis. As compared to the good clustering performance from SAUCIE embedding (**Supplementary Note 1 Fig. 5b)**, these two DR methods' embeddings fail to cluster tumor cells in one cluster (red and green clusters). Further, KPCA RBF's blue and purple cells are noticeably mixed together as well. Source data are provided as a Source Data file.

#### Supplementary Discussion 1: Discussion of Scalability and Dimensionality

As discussed in the **Backgrounds** section of our paper, a unique differentiating factor between CyTOF and scRNA-seq is the dimensionality of the datasets. Namely, CyTOF has orders of magnitude less channels (features denoted as $P$) and much more cells (sample size denoted as $N$ ). The scalability benchmark clearly indicates that different methods scale vastly differently with respect to both $N$ but not so much for $P$. Here, we discuss efficiency concerns of some of the prominent methods and their fit for CyTOF data analysis purposes.

##### SAUCIE

SAUCIE[1] is based on an autoencoder implemented in tensorflow. It uses an 8-layer neural network, and DR is achieved by extracting the middle bottleneck layer. While SAUCIE does not give a specific efficiency claim in terms of N or P, both its author and our study found that its efficiency for DR is top-tier. Surprisingly, the performance of SAUCIE, even without using GPUs, is on par with the most efficient methods tested. Given the nature of the autoencoder, only the input and output layers have to have the same size as P of the original training data. Thus, the impact of  P should be small, unless larger intermediate layers are needed for higher dimensional data like scRNA-seq. Sample size N can potentially interact with the choice of batch size which may in turn impact scalability, but our results suggest that default settings already produce good embeddings for CyTOF. From our study, SAUCIE remains fast and accurate for CyTOF samples. Thus, CyTOF's dimensionality is not of concern for SAUCIE.

##### scvis

scvis[2] uses variational inference to find the latent variable to represent the latent variable representing underlying structure in the data. To aid the visualization of the latent space, it utilizes the idea of tSNE as a regularizer: specifically, it uses KL divergence to minimize distance as defined by a Gaussian kernel. While scvis is among the least efficient methods without memory bounds, we found that the number of features does not play a role in its runtime with typical CyTOF feature spaces. However, sample size $N$ poses a significant concern here since a typical sample can take as much as 12 hours to run on a server even though runtime does tend to plateau. The use of scvis is thus a compromise between theoretical fit and practicality.

##### UMAP

UPAM falls under the second tier of methods in terms of efficiency. In UMAP's methodological paper[3], the authors claimed approximately $O(N^{1.14})$ for the optimization algorithm. Since UMAP falls in the same category of algorithms that utilize k-nearest neighbors graphs, its performance mostly depends on sample size $P$ rather than $N$ like Fit-SNE. However, due to the fact that it does not need to normalize the density in the optimization step, the authors claimed that UMAP can be much more efficient when reducing dimensions down to more than 2 or 3 dimensions. While higher dimensional DR embeddings are not the focus of this paper, it is a common workflow in scRNA-seq data analysis, which may help explain the advantage of UMAP in that setting.

**PHATE**

PHATE[4] is a method that focuses the differentiation path of cells. Namely, it calculates the local similarity and global relationship. Then, it uses potential distance to account for the overall structure and metric MDS for visualization. There is no explicit claim of efficiency, but we observed that it is among the least efficient methods for runtime. Like other methods, dimensionality does not play a large role for efficiency except for using larger vectors, but it scales at least quadratically with the number of cells. Like scvis, the original paper also used PHATE for scRNA-seq data. Therefore, dimensionality should not be much of a concern given that it is a distance-based method, but sample size again poses a challenge.

**SQuaD-MDS**

SQuaD-MDS[5] improves upon the original MDS by employing Stochastic Quartet Descent. CyTOF's large sample size $N$ renders MDS unusable without downsampling because of $O(N^2)$ memory complexity. SQuaD-MDS vastly improves the efficiency by reducing runtime and memory complexity down to $O(N)$, which is on par with the second tier of methods. As with other force-directed methods, dimensionality does not play nearly as large of a role as $N$. SQuaD-MDS is one of the methods that is general-purpose. However, it was tested on both image data with high dimensions and some low-dimensional scRNA-seq benchmark datasets by its authors. We can thus reasonably expect that SQuaD-MDS will work for CyTOF and scRNA-seq datasets across a wide range of dimensionality.

**tSNE (All major variants)**

As tested extensively in this study, tSNE's implementation has important ramification in terms of scalability. While we included two implementations in main benchmarks and five in **Sup. Fig. 4**, these implementations can be divided into the following categories:

1. Original tSNE[6]
2. Barnes-Hut tSNE[7]
3. Fit-SNE[8]

The original tSNE has $O(N^2)$ for computation time and memory efficiency. This is obviously unrealistic for large CyTOF samples: in fact, any samples with $N > 10,000$ can start to become problematic. As tSNE's optimization involves the pairwise forces exerted by each point in both the original and DR space, the main bottleneck is $N$ rather than $P$.

The Barnes-Hut optimization for tSNE (BH tSNE) reduces the run time to $O(N\ logN)$ for computation and $O(N)$ for memory usage. In our benchmarks, BH tSNE can be realistically used for most CyTOF datasets, but significant time is needed when samples are large. Recently, Fit-SNE further reduces runtime to $O(2pN)$. Fit-SNE is by far the fastest implementation we tested while also being the most competitive with other methods (*e.g.* UMAP). The one caveat with Fit-SNE is its availability since it is still relatively new, but there are already implementations and wrappers available for use.
