## Supplementary Data Legends for "Comparative Analysis of Dimension Reduction Methods for Cytometry by Time-of-Flight Data"

**Supplementary Data 1 All DR methods evaluated in this work.** Basic information on each DR method and their usability criteria are listed.

**Supplementary Data 2 CyTOF and scRNA-seq datasets used in this study.** The data include metadata on each cohort along with links to their publication and repositories.

**Supplementary Data 3 Explanation of the DR validation metrics employed in this study.** For each evaluation metric, its implementation, references, weight, and any additional notes are included. Green cells correspond to accuracy metrics, whose weights are used to calculate the main ranking of methods, and blue cells are for other metrics.

**Supplementary Data 4 Tuning parameters tested for the DR methods.** For each method tuned, all tuned parameters are listed along with their optimal settings. Note that all tuning parameters in other benchmarks, other than the parameter tuning benchmark, use the software implementations' default. See Methods section for details.
